# Enhanced Effector Activity of Mediator Kinase Module Deficient CAR-T Cells

**DOI:** 10.1101/2022.09.02.506235

**Authors:** Katherine A. Freitas, Julia A. Belk, Elena Sotillo, Bence Daniel, Katalin Sandor, Dorota Klysz, Vandon T. Duong, Kylie Burdsall, Peng Xu, Meena Malipatlolla, Micah G. Donovan, Evan W. Weber, Robbie G. Majzner, Howard Y. Chang, Joaquin M. Espinosa, Ansuman T. Satpathy, Crystal L. Mackall

## Abstract

Adoptive T cell immune therapies mediate impressive clinical benefit in a fraction of patients, but anti-tumor effects are often limited by inadequate T cell potency. To identify genes limiting T cell effector function, we conducted genome-wide CRISPR knock-out screens in human primary CAR-T cells. The top hits were *MED12* and *CCNC*, components of the cyclin-dependent kinase (CDK) module of the Mediator complex, an evolutionarily conserved regulator of gene transcription. *MED12* or *CCNC* deficient CAR-T cells manifest increased expansion, cytokine production, metabolic fitness, effector function, anti-tumor activity and reduced terminal effector differentiation. Chemical inhibition of CDK8/19 kinase activity recapitulated some features of genetic loss of *MED12*, including increased T cell expansion. *MED12* deficient CAR-T cells showed widespread but selective increases in chromatin accessibility, MED1 chromatin occupancy, and H3K27 acetylation at enhancers used by transcription factors playing a critical role in T cell fate, including several STAT and AP1 family members. The most pronounced enhancement was observed for STAT5 which manifested as increased sensitivity to IL-2 in *MED12* deficient T cells. These results link Mediator induced transcriptional coactivation with T cell effector programming and identify the CDK module as a target for enhancing the potency of anti-tumor T cell responses.

**One Sentence Summary:** The Mediator kinase module is a primary regulator of T cell differentiation, and genetic or small molecule-based inhibition of this module enhances effector T cell potency.

## Introduction

T cell based immunotherapy including both immune checkpoint inhibition and adoptive cell therapies have demonstrated impressive activity for the treatment of many cancers^1–8^, but durable responses are not achieved in most patients. A central barrier to progress is limited T cell potency, resulting from a myriad of factors including T cell exhaustion, senescence, anergy and local and systemic immunosuppression^9–12^. Advances in understanding the biology of T cell exhaustion are providing novel approaches to prevent this phenomena, including overexpression of c-JUN^13^, deletion of NR4A^14^, and transient induction of T cell rest, which can reverse the epigenetic imprint of exhaustion^15^. However, it remains unclear whether exhaustion resistance will be sufficient to overcome the multitude of immunosuppressive factors within the tumor microenvironment.

CRISPR technology provides unparalleled opportunities to engineer human cells. *Ex vivo* CRISPR has already been used safely to deliver gene-edited tumor-specific T cells to humans with cancer^16^, and *in vivo* CRISPR mediated gene editing was recently demonstrated^17^. The CRISPR platform has also been optimized to conduct forward genetic screens in primary human T cells to identify novel targets to augment T cell function,^18^ including genes that regulate PD-1 expression^19^, T cell proliferation, and resistance to adenosine-mediated immunosuppression^18^. Thus, advances in gene editing are providing accessible approaches to deliver tumor-specific T cells programmed for enhanced potency, but much work remains to be done to define the specific genes for which editing will most potently augment anti-tumor responses in humans.

We utilized CRISPR screening to identify genes that regulate effector function in primary human T cells expressing chimeric antigen receptors (CARs) and discovered that *MED12* and *CCNC*, genes encoding proteins in the CDK module of the Mediator complex, negatively regulate T cell effector differentiation. Mediator, an evolutionarily conserved multi-subunit protein complex that acts as a bridge between enhancer-bound transcription factors and the general transcription machinery^20^, is required for gene transcription and plays a central role in establishing cellular identity by coordinating transcriptional networks^21,22^. Across multiple CAR-T cell models with different costimulatory domains, we observed that genetic disruption of the CDK module of Mediator induced widespread transcriptional and epigenetic changes that resulted in enhanced effector function and metabolic fitness, diminished terminal effector differentiation, and increased anti-tumor activity, and small molecule mediated inhibition of CDK8 phenocopied several of these enhancements. These results implicate the Mediator CDK module as a therapeutic target for augmenting T cell function and identify a previously unknown role for *MED12* in regulating human T cell differentiation.

## Results

### Genome-wide screen identifies the Mediator CDK module as a regulator of CAR-T cell expansion and cytokine production

To identify genes which restrain CAR-T cell function in the setting of T cell exhaustion, we performed two genome-wide CRISPR deletion screens in human primary T cells expressing HA-28ζ, a high affinity GD2-targeting CAR that induces functional, transcriptomic and epigenetic hallmarks of T cell exhaustion via tonic signaling, thereby mimicking chronic exposure to antigen^13,15^. Using a previously published sgRNA library^23^, editing was achieved by adapting the SLICE platform (single guide RNA (sgRNA) lentiviral infection with Cas9 protein electroporation)^18^ to incorporate CAR transduction (**Fig. 1A and S1A**). We detected 98% of the sgRNA library in transduced CAR-T cells, with 19,885 genes targeted by at least 4 sgRNAs (**Fig. S1B-C**). Successful editing was confirmed by drop out of sgRNAs targeting a “gold standard” set of essential genes but not control guides after 23 days in culture (**Fig. S1D, Table S1, Table S2**).

**Figure 1.**
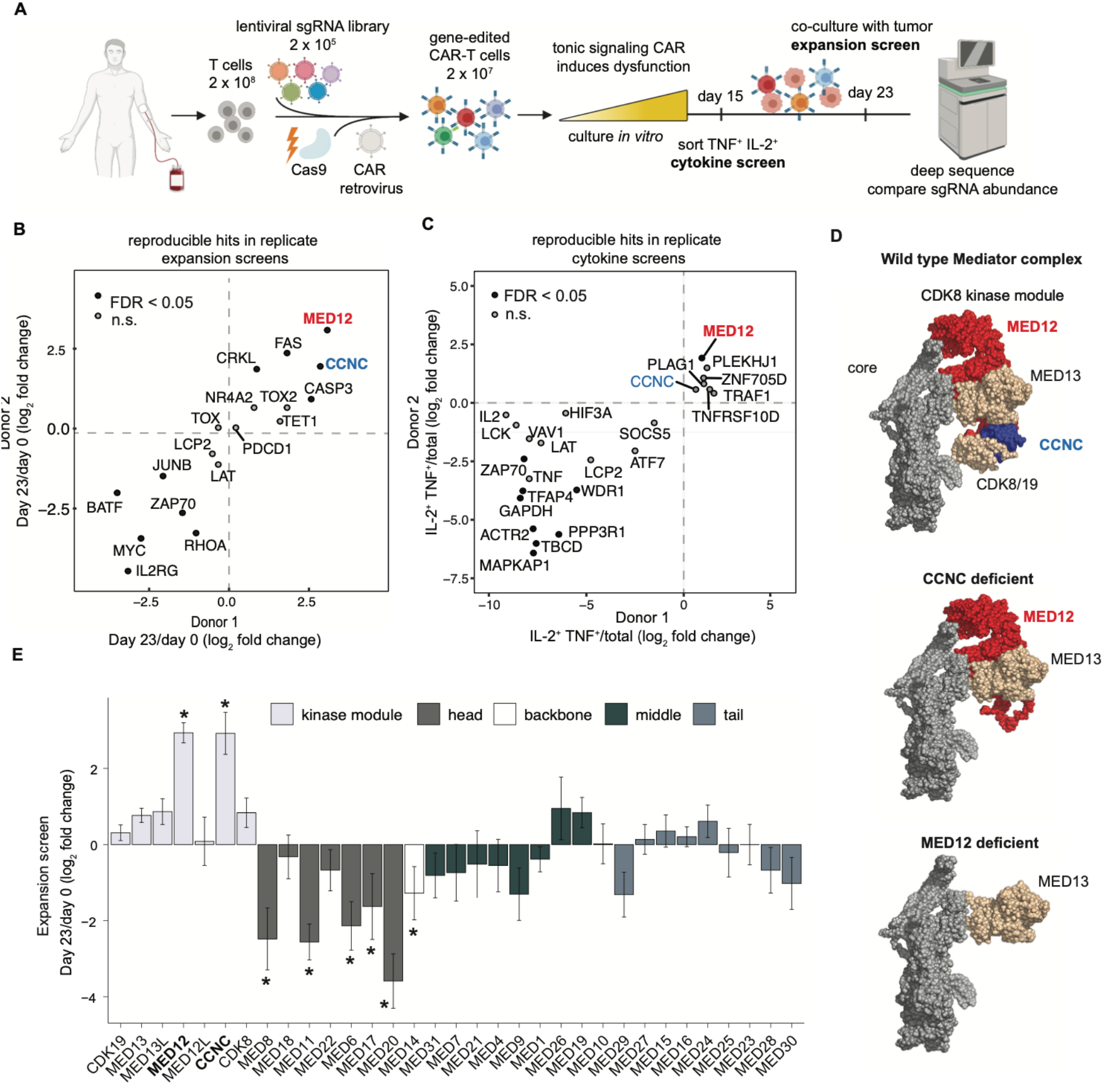
Genome-wide CRISPR screen identifies subunits of the Mediator kinase module as regulators of CAR-T cell effector function. **A)** Schematic depicting CRISPR knockout screen for regulators of cytokine production and CAR-T cell expansion using a tonic signaling model of CAR-T cell exhaustion. **B)** Enrichment of gene knockouts in replicate expansion screens. CRISPR-edited HA-28ζ CAR-T cells were generated from two donors, cultured *in vitro* for 15 days, and then co-cultured with GD2^+^ tumor cells until day 23. **C)** Enrichment of gene knockouts in replicate cytokine production screens. CRISPR-edited HA-28ζ CAR-T cells were generated from two donors, cultured *in vitro* for 15 days, stimulated with GD2^+^ tumor cells, and the top 10% of TNFα and IL-2 expressing cells were isolated by FACS. **D)** Predicted Cryo-EM structure of yeast Mediator complex (top) showing the effect that depletion of Cyclin C (*CCNC* deficient) or *MED12* (*MED12* deficient) would have on assembly of other subunits. Core Mediator is shown in grey, CDK8 kinase module is colored. Representations were created with Chimera using Protein Data Bank accessions 7KPX and 5U0P. **E)** Bar graphs depicting enrichment of sgRNA targeting all Mediator complex subunits in the expansion screen. Data are mean ± s.d. (*n* = 4-10 guides per genes). Colors indicate module of the Mediator complex assigned to each subunit. **B, C, E)** Data are mean of *n* = 4-10 guides per gene. Data are pooled from two independent experiments (*n* = 2 donors). Gene-level statistical significance was determined by the MAGeCK algorithm. *\*FDR* < 0.05.

Replicate screens were conducted using HA-28ζ CAR-T cells from two donors to identify negative regulators of T cell expansion and cytokine production, both critical elements in the effector T cell response. For the expansion screen, we cultured the transduced cells *in vitro* for 15 days, then co-cultured with GD2^+^ tumor cells until day 23 and compared sgRNA abundance between day 0 and 23 (**Fig. S1E**). Using the MAGeCK algorithm^24^ both donors showed enrichment of sgRNAs targeting genes known to inhibit T cell survival, such as *FAS* and *CASP3*^25^ (**Fig. 1B**), while sgRNAs targeting genes known to promote T cell proliferation, such as *IL2RG, MYC* and *ZAP70*, were depleted. The expansion screen also showed depletion of sgRNAs targeting *BATF* and *JUNB*, suggesting a survival role for AP-1 family members in the setting of chronic stimulation. The top hits in the expansion screen were *CCNC* and *MED12*, members of the CDK module of Mediator, with 7 out of 7 guides targeting *CCNC* and 8 out of 8 guides targeting *MED12* positively enriched **(Fig. S2-C**).

In a second screen we sought to identify genes restraining antigen-induced cytokine production by culturing HA-28ζ CAR-T cell knock-out libraries *in vitro* for 15 days, adding GD2^+^ tumor cells to the co-culture for 6 hours, then using FACS to sort cells and compare sgRNA abundance in CAR-T cells expressing IL-2 and TNFα protein against the total population (**Fig. S1E)**. *MED12* was also enriched in the cytokine screen, with 6 out of 7 guides demonstrating enrichment, while several genes, including 9 out of 9 guides targeting *ZAP70* were depleted, consistent with the known role for *ZAP70* in CAR signaling^26^ (**Fig. 1C, Fig. S1B-C**). Six of 9 guides targeting *TNF* and 5 out of 8 guides targeting *IL2* were significantly depleted, demonstrating an expected loss of cytokine-negative cells in the sorted population (**Fig. S2B**).

Mediator consists of a 26 subunit core organized into head, middle, backbone, and tail domains, and a 4 subunit dissociable kinase module^27,28^ (**Fig. 1D**). sgRNAs targeting all members of the kinase module were positively enriched in the expansion screen except for *MED12L*, which is not expressed in T cells (**Fig. 1E**). *CCNC* and *MED12* are both centrally located in the kinase module^29^, suggesting that loss of either gene disrupts a common function. In contrast, sgRNAs targeting subunits of the head, backbone, and middle domains of core Mediator were associated with poor expansion **(Fig. 1E**). Together, in this model utilizing CAR-T cells rendered dysfunctional due to T cell exhaustion, we demonstrated a requirement for the core Mediator complex in T cell survival and a regulatory role for the Mediator kinase module in T cell expansion and cytokine production.

### MED12 deficient and CCNC deficient exhausted and non-exhausted CAR-T cells demonstrate increased in vitro and in vivo expansion independent of co-stimulation domain

To validate the expansion screen findings, sgRNAs targeting *CCNC, MED12*, or *AAVS1* as a control, were delivered as ribonucleoprotein 3 days after T cell activation followed by retroviral transduction of the HA-28ζ CAR. *CCNC* and *MED12* deletion were confirmed by immunoblotting and Sanger sequencing using Inference of CRISPR Edits (ICE)^30^ (**Fig. S3A-B**). *MED12* and *CCNC* deficient HA-28ζ CAR-T cells showed greater expansion compared to control cells over 23 days in culture (**Fig. 2A)** and following serial stimulation with tumor cells (**Fig. S3C)**. Because CAR transduction efficiency, as well as the ratio of CD4^+^ to CD8^+^ cells, could impact CAR-T cell function, we confirmed loss of *MED12* or *CCNC* did not change CAR expression or the ratio of CD4^+^ to CD8^+^ cells (**Fig. S3D-F**). Together, these results confirm that *MED12* and *CCNC* deficient human HA-28ζ CAR-T cells manifest enhanced antigen-driven expansion.

**Figure 2.**
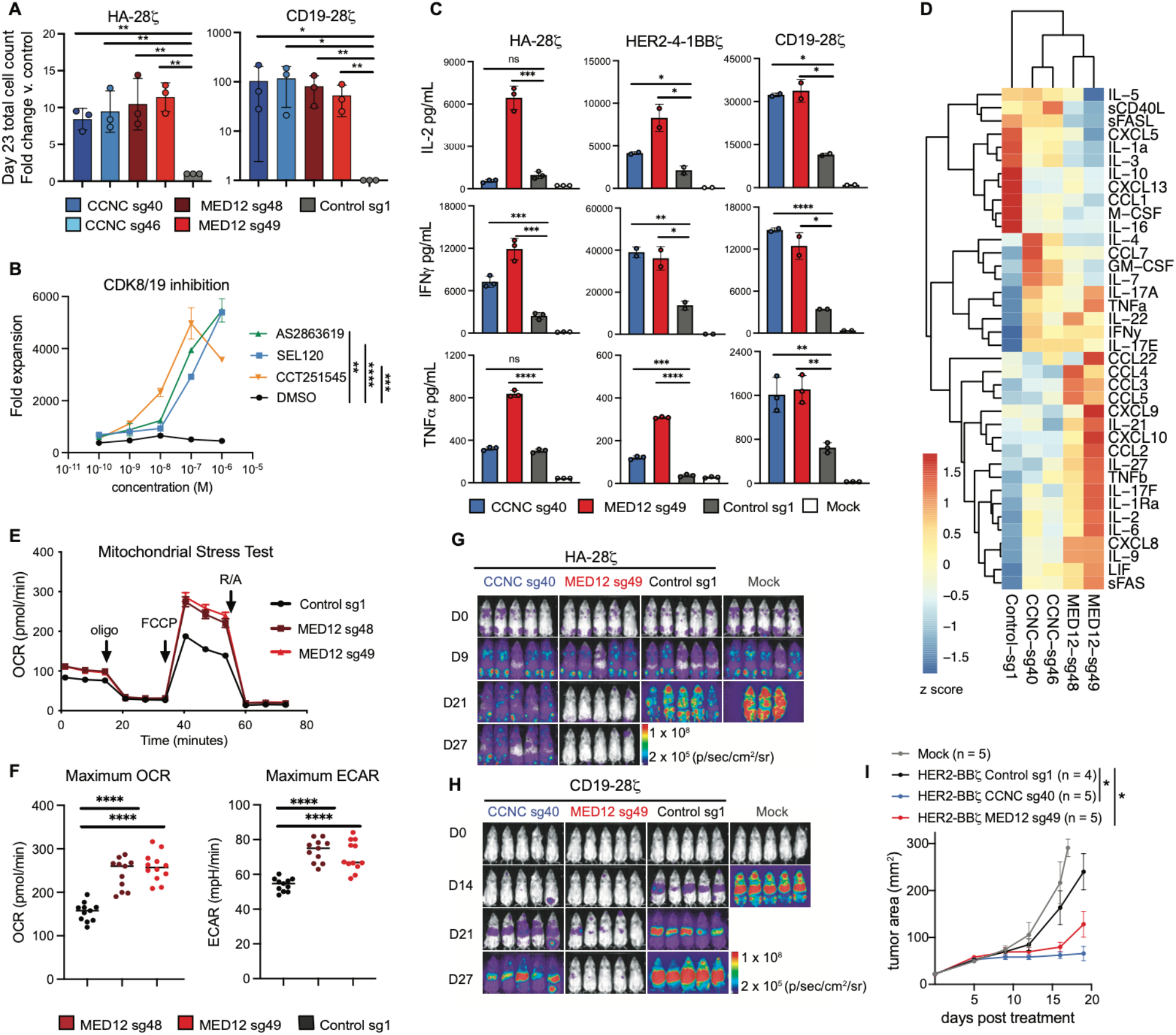
Disruption of Mediator kinase module in CAR-T cells increases effector function indicated by T cell expansion, cytokine production, metabolic rate, and tumor clearance. **A)** *In vitro* expansion of *CCNC* and *MED12* deficient HA-28ζ (left) and CD19-28ζ (right) CAR-T cells. Fold change in total cell count after 23 days in culture relative to control CAR-T cells edited at the safe harbor AAVS1 locus. Two unique sgRNAs were used to validate each candidate gene. Data are mean ± s.d. of *n* = 3 donors. Ratio paired *t-* test. \**P* < 0.05, \*\**P* < 0.01, \*\*\**P* < 0.001, \*\*\*\**P* < 0.0001 **B)** *In vitro* expansion of human primary T cells with duals inhibitors of CDK8 and CDK19 over 15 days in culture. Inhibitors were supplemented to the media every 48 hours. *n* = 2 wells. Representative of 3 independent experiments. **C)** IL-2 (top), IFNγ (middle), and TNFα (bottom) release after 24-hour co-culture with tumor cells as measured by ELISA. Mock or *CCNC* or *MED12* deficient HA-28ζ, CD19-28ζ and HER2-4-1BBζ CAR-T cells were stimulated 1:1 with NALM6-GD2, NALM6, or 143B cells respectively. Data are mean ± s.d. from duplicate or triplicate wells. Representative results of *n* = 4 donors (HA-28ζ, CD19-28ζ) or *n* = 2 donors (HER2-4-1BBζ). **D)** Heatmap of 38 cytokines produced by *CCNC* or *MED12* deficient CD19-28ζ CAR-T cells following 24-hour co-culture with NALM6 leukemia cells. Data are mean from duplicate wells in a multiplex bead-based assay. Two unique sgRNAs were used to validate each candidate gene. **E and F)** Seahorse analysis of oxygen consumption rate (OCR) and extracellular acidification rate (ECAR) of control or *MED12* deficient CD19-28ζ CAR-T cells under resting and challenge conditions. Data are mean of *n* = 12 replicate wells. Representative results from two independent experiments. **G)** NSG mice were injected intravenously with 1.0 ×10^6^ NALM6-GD2-Luc leukemia cells and treated with 2.0 × 10^5^ mock or *CCNC* or *MED12* deficient HA-28ζ CAR-T cells 9 days after tumor infusion (*n* = 5 mice). **H)** NSG mice were injected intravenously with 1.0 ×10^6^ NALM6-Luc leukemia and treated with 2.5 × 10^5^ mock or *CCNC* or *MED12* deficient CD19-28ζ CAR-T cells 3 days after tumor infusion (*n* = 5 mice). **I)** Tumor area of NSG mice injected intramuscular with 1 × 10^6^ 143B osteosarcoma cells and treated 4 days later with 5 × 10^6^ mock or *CCNC* or *MED12* deficient HER2-4-1BBζ CAR-T cells. Tumor area was measured by caliper. Two-way ANOVA test with Dunnett’s multiple comparison test. \**P* < 0.01. **G, H, and I**) Representative experiment from two independent experiments (*n* = 2 donors). **B, C and F)** Two-tailed unpaired Student’s *t*-test. \**P* < 0.05, \*\**P* < 0.01, \*\*\**P* < 0.001, \*\*\*\**P* < 0.0001

To determine whether the findings were specific for exhausted T cells or resulted from a more generalized enhancement in T cell proliferative capacity, we deleted *MED12* and *CCNC* in T cells expressing the CD19-28ζ CAR, which do not acquire features of exhaustion when cultured *in vitro. CCNC* and *MED12* deficient CD19-28ζ CAR-T cells also demonstrated increased expansion *in vitro* and adoptively transferred *MED12* and *CCNC* deficient HA-28ζ and CD19-28ζ CAR-T cells showed increased *in vivo* expansion in tumor-bearing NSG mice compared with control CAR-T cells (**Fig. 2A, Fig. S3G)**. *MED12* and *CCNC* behave as tumor suppressors in some settings^31,32^, however when we removed IL-2 from the culture medium, we observed a complete loss of viable cells within 3 weeks (**Fig. S3H**), indicating that expansion is not associated with transformation as the cells remain IL-2 dependent. We also observed enhanced expansion of *MED12* and *CCNC* deficient HER2-4-1BBζ CAR-T cells (**Fig. S3F**), confirming that the findings are not restricted to CAR-T cells incorporating a CD28 costimulatory domain. To determine if these effects were dependent on catalytic activity of the CDK module, we cultured T cells with compounds that are dual inhibitors of CDK8 and CDK19 and observed significant increases in T cell expansion (**Fig. 2B**). Together, these results demonstrate that perturbation of the CDK module increases expansion in both exhausted and non-exhausted CAR-T cells regardless of whether the CAR incorporates a CD28 or 4-1BB co-stimulatory domain.

### CCNC deficient and MED12 deficient CAR-T cells produce higher levels of inflammatory cytokines following antigen stimulation

Next, we sought to confirm the effects of *MED12* and *CCNC* deletion on antigen-induced cytokine production using bulk assays and by assessing single cell production of IL-2 and TNFα by flow cytometry. Bulk cultures of antigen stimulated *MED12* and *CCNC* deficient HA-28ζ, CD19-28ζ and HER2-4-1BBζ CAR-T cells produced higher levels of cytokines **(Fig. 2C)** and *MED12* and *CCNC* deficient HA-28ζ and CD19-28ζ CAR-T cells demonstrated increased frequencies of IL-2 and TNFα expressing cells (**Fig. S4A-B**). Additionally, we found elevated *IL2, IFNG*, and *TNF* mRNA levels in *MED12* deficient cells compared with control cells, suggesting these changes are transcriptionally mediated (**Fig. S4C**). To assess the impact of Mediator kinase module disruption more broadly on antigen-induced cytokine secretion, we performed bead-based multiplex immunoassay profiling of 38 cytokines in supernatants collected from CD19-28ζ CAR-T stimulated with CD19^+^ NALM6 leukemia cells for 24 hours. Hierarchical clustering showed that the cytokine profile of *MED12* and *CCNC* deficient CD19-28ζ CAR-T cells was distinct from controls (**Fig. 2D**), with increased proinflammatory cytokines including IFNγ, TNFα, IL-17, IL-6, increased inflammatory chemokines CXCL10 and CCL3, and increased common gamma chain family cytokines IL-2 and IL-9, which promote T cell survival and differentiation (**Fig. S4D**). In contrast, *MED12* and *CCNC* deficient cells produced lower levels of the immunosuppressive cytokine IL-10. Together, these results demonstrate that *MED12* and *CCNC* constrain antigen induced T cell expansion and inflammatory cytokine production, while promoting IL-10 production, and raise the prospect that *MED12* or *CCNC* deficient T cells may demonstrate enhanced anti-tumor immune responses.

### CCNC deficient and MED12 deficient CAR-T cells demonstrate increased cytotoxicity, metabolic fitness, and anti-tumor activity

*MED12* or *CCNC* deficient HA-28ζ CAR-T cells demonstrated increased cytotoxicity *in vitro* following repeated stimulation with NALM6-GD2 leukemia cells (**Fig. S5A-B**). To further assess whether *MED12* or *CCNC* deficient T cells manifest metabolic features of enhanced effector functionality^33^, we measured glycolytic rates and basal and maximal oxygen consumption rates. We observed increased basal and maximal oxygen consumption in *MED12* and *CCNC* deficient cells, despite no change in mitochondrial mass, and increased basal and maximal rates of glycolysis (**Fig. 2E-F, Fig. S5C-D)**. *CCNC* and *MED12* deficient CAR-T cells also demonstrated enhanced anti-tumor function *in vivo*, in a model wherein NSG mice were inoculated with NALM6 or NALM6-GD2 leukemic cells expressing luciferase and treated 3 days later with gene-edited CD19-28ζ or HA-28ζ CAR-T cells respectively. Using bioluminescence imaging to track tumor progression, we observed increased tumor clearance and prolonged survival in mice treated with *MED12* and *CCNC* deficient CAR-T cells compared to the control group (**Fig. 2G-H, Fig. S5E-F**). Similarly, we observed enhanced tumor control and prolonged survival in mice engrafted with 143B osteosarcoma cells and treated with *MED12* and *CCNC* deficient CAR-T cells expressing the HER2-4-1BBζ receptor (**Fig. 2I, Fig. S5G**). Collectively, these results demonstrate that *MED12* and *CCNC* deficient T cells manifest enhanced hallmark features of effector cells, spanning antigen induced expansion, cytokine production, metabolic fitness and killing capacity.

### MED12 deficient CAR-T cells have an effector-like phenotype with reduced terminal differentiation

To more deeply characterize the impact of Mediator CDK module disruption on T cell differentiation, we focused on *MED12* because this gene was a common hit in both the expansion and cytokine screens. *MED12* deficient CD19-28ζ CAR-T cells were cultured *in vitro* until day 23 and then analyzed for CCR7 and CD45RO expression to quantify stem cell memory, central memory, effector memory, and terminal effector subsets. Both control and *MED12* deficient CAR-T cells displayed effector phenotypes based upon an absence of CCR7; however, *MED12* deficient CAR-T cells expressed high levels of CD45RO, while control cells were largely negative for this marker at this timepoint, suggesting that loss of *MED12* may prevent terminal differentiation of effector cells (**Fig. 3A-B**)^34–36^. To further characterize the *MED12* deficient phenotype, we undertook single cell proteomic analysis of 34 proteins using mass cytometry, to measure lineage-defining transcription factors and cell surface markers associated with activation, exhaustion, and T cell differentiation (**Table S3, Fig. S6A**). Both *MED12* deficient and control CD19-28ζ CAR-T cells expressed high levels of Blimp-1 and low levels of Eomes and CD28, consistent with an effector phenotype^37^, however, unbiased clustering demonstrated significant distinctions between *MED12* deficient and control phenotypes (**Fig. 3C, Fig. S6B**). *MED12* deficient cells expressed higher levels of T-bet, TOX, and CD45RO, and lower levels of CD45RA and IL7R, a phenotype associated with short lived effector cells (SLECs) that have not undergone terminal differentiation^38^ (**Fig. 3D, Fig. S6C-D**). Paradoxically however, *MED12* deficient cells also manifested high levels of CD62L, which is usually associated with stem cell and central memory subsets and not typically expressed by SLECs. *MED12* deficient CD19-28ζ CAR-T cells also expressed high levels of LAG-3, while other exhaustion markers PD-1, TIM3, TIGIT, and CD39 were lowly expressed and unchanged from control cells **(Fig. 3D, Fig. S6E)**. *MED12* deficient HA-28ζ CAR-T cells also expressed high levels of LAG-3, modest but statistically significant increases in TIM3 and PD-1, but diminished levels of CD39, a marker that is associated with terminal exhaustion and diminished stemness^39,40^ (**Fig. 3E**). Together, these results demonstrate that *MED12* deficient CAR-T cells manifest expansion of a unique CCR7^−^IL7R^−^Tbet^+^CD62L^+^ effector cell subset that displays enhanced cytokine production, effector cell potency, metabolic fitness, and antitumor activity.

**Figure 3.**
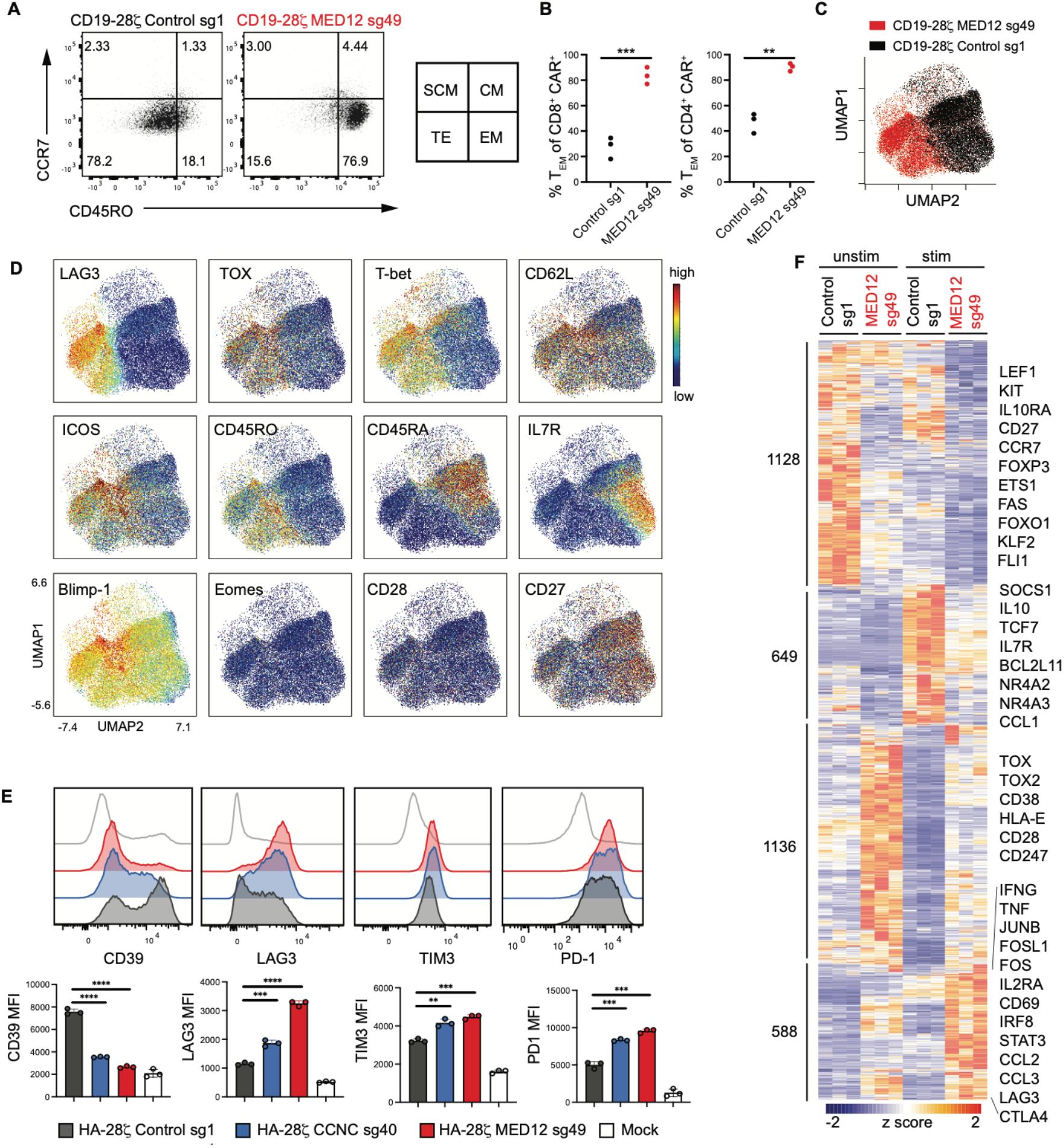
*MED12* deficient CD19-28ζ CAR-T cells have an effector-like phenotype with reduced terminal effector differentiation. **A)** Flow cytometry analysis of T cell subsets as assessed by CD45RO and CCR7 expression in control or *MED12* deficient CD8^+^ CD19-28ζ CAR-T cells 23 days after T cell activation. Representative result of *n* = 3 donors. Gating and subtyping strategy shown (SCM, Stem Central Memory; CM, Central Memory, TE, Terminal Effector; EM, Effector Memory) **B)** Frequency of T effector memory cells (CD45RO^+^, CCR7^-^) in CD8^+^ and CD4^+^ CAR-T cells. *n* = 3 donors. Two-tailed paired Student’s *t*-test. \*\**P* < 0.01, \*\*\**P* < 0.001. **C and D)** Uniform Manifold Approximation and Projection (UMAP) analysis of control and *MED12* deficient CD19-28ζ CAR-T cells 15 days after T cell activation. Expression of 34 markers was analyzed by CyTOF. Representative donor of *n* = 3 donors. Control and *MED12* deficient samples are combined and colored **C)** by genotype or **D)** by marker intensity. **E)** Flow cytometry histograms (top) and quantification (bottom) of exhaustion markers in *CCNC* and *MED12* deficient HA-28ζ CAR-T cells. Data are mean and s.d. of *n* = 3 wells. Representative result of two independent experiments. Two-tailed unpaired Student’s *t*-test. \**P* < 0.05, \*\**P* < 0.01, \*\*\**P* < 0.001, \*\*\*\**P* < 0.0001. **F)** Heatmap of differentially expressed genes in control or *MED12* deficient CD19-28ζ CAR-T cells detected by bulk RNA-seq 15 days after T cell activation. Cells were collected on day 15 (unstim) or stimulated (stim) through the CAR for 3 hours with plate-bound anti-idiotype antibody. Adjusted *P* < 0.01. *n* = 3 donors.

To assess genome-wide transcriptional differences between *MED12* deficient and control cells, bulk RNA-seq was performed on day 15 CD19-28ζ CAR-T cells following a 3-hour stimulation via the CAR *in vitro*. Principal component analysis (PCA) demonstrated that *MED12* deficient were transcriptionally distinct from control cells (**Fig. S7A)** with differential expression of 3501 genes between genotypes in at least one condition (**Fig. 3F**). Consistent with the functional and phenotypic data, *MED12* deficient cells demonstrated increased expression of numerous genes associated with effector cell differentiation, including AP-1 family transcription factors (*FOS, JUNB, BATF, BATF3*), *IFNG, TNF, CD38, IL2RA*, and *CD69*. They also expressed lower levels of genes associated with T cell stemness including *LEF1, TCF7, CD27, IL7R*, and *KLF2* and decreased expression of several genes associated with immune suppression including *FOXP3, IL-10*, and *IL-10R*. Together, the results support a model wherein *MED12* promotes terminal differentiation of effector T cells while *MED12* deficient T cells manifest sustained effector programming and reduced terminal differentiation.

### Loss of MED12 increases chromatin accessibility in CD19-28ζ CAR-T cells and promotes T effector transcriptional programming

The Mediator complex can regulate chromatin architecture through DNA looping^41^, thus disruption of the complex could have broad effects on the chromatin landscape. To test this, we assessed chromatin accessibility using Assay for Transposase Accessible Chromatin (ATAC-seq) in resting and stimulated *MED12* deficient and control CD19-28*ζ* CAR-T cells. PCA showed 40.5% of the variance (PC1) was attributed to the effect of stimulation, while PC2 (24% of the variance) could be attributed to loss of *MED12* (**Fig. 4A**), with similar separation by PC2 in both unstimulated and stimulated conditions. These results demonstrate that the changes induced by *MED12* deletion are largely independent of stimulation. We observed increased accessibility of 4670 peaks and decreased accessibility of 1927 peaks in resting *MED12* deficient cells, and increased accessibility of 6769 peaks and decreased accessibility of 1058 peaks in stimulated *MED12* deficient cells, consistent with *MED12* deficiency inducing a pattern of increased chromatin accessibility (**Fig. 4B**). As expected, T cell stimulation preferentially opened chromatin, but this effect was even more pronounced in *MED12* deficient cells (**Fig. S7B**), as k-means clustering revealed several regions that were moderately more accessible in *MED12* deficient versus control CAR-T cells under resting conditions, but dramatically more accessible upon stimulation (**Fig. 4C**). The chromatin accessibility changes revealed by ATAC-seq in *MED12* deficient cells correlated with gene expression (**Fig. 4D**).

**Figure 4.**
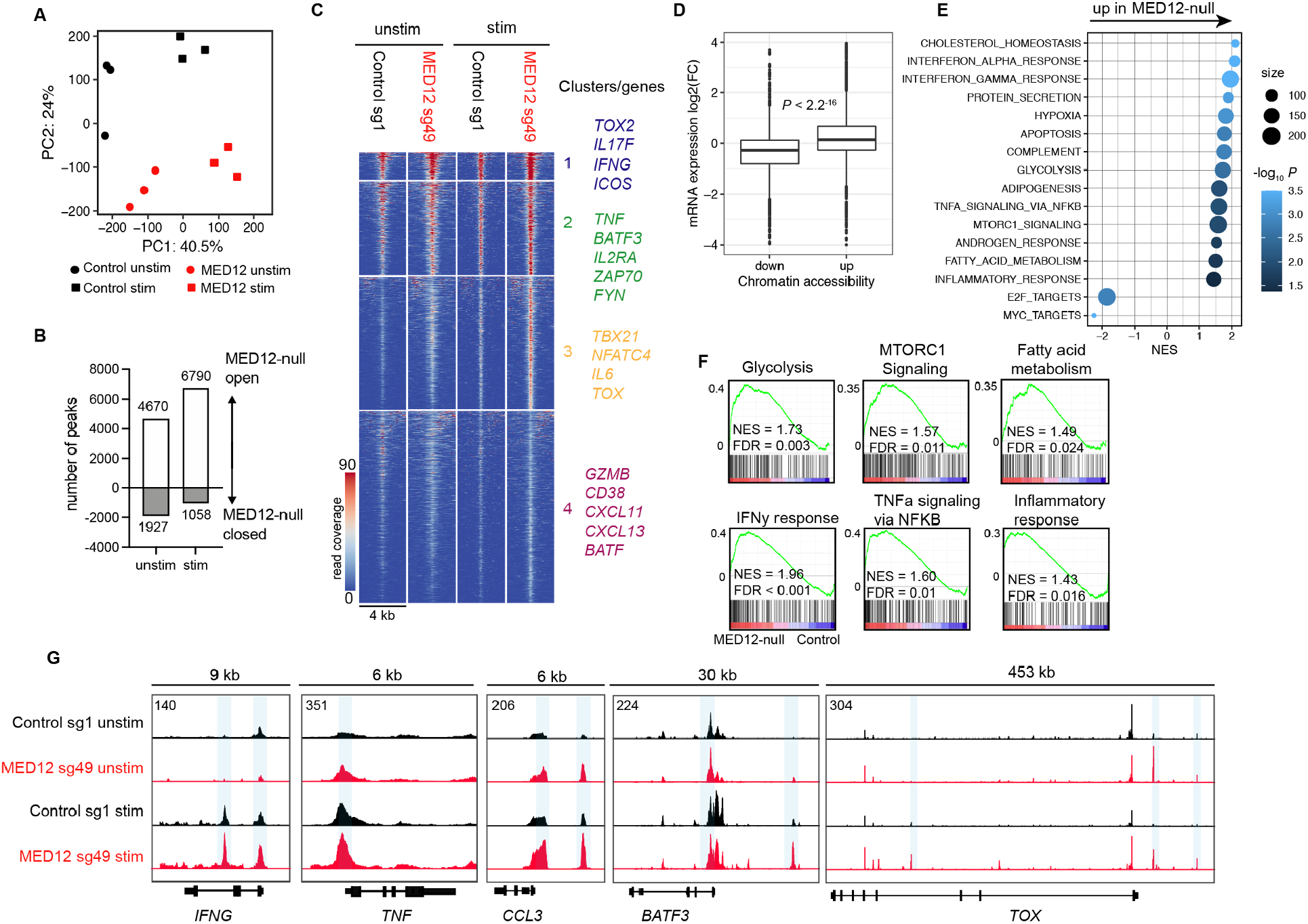
Loss of *MED12* increases chromatin accessibility in CD19-28ζ CAR-T cells and promotes T effector transcriptional programming. **A)** PCA of ATAC-seq performed on control and *MED12* deficient CD19-28ζ CAR-T cells 15 days after T cell activation. Cells were collected without stimulation (unstim) or stimulated (stim) for three hours with plate-bound anti-idiotype antibody. **B)** Number of peaks with significant change in accessibility detected by ATAC-seq between *MED12* deficient and control cells. Log2FC > 1 and adjusted *P* < 0.05. **C)** Chromatin accessibility of control and *MED12* deficient CD19-28ζ CAR-T cells. Each row represents one peak displayed over a 4-kb window. Log2FC > 1 and adjusted *P* < 0.05. Regions were clustered with k-means clustering. **D)** Boxplot depicting fold change in mRNA expression between unstimulated *MED12* deficient and control CAR-T cells at genes proximal to OCRs with differential chromatin accessibility. Kruskal Wallis test. **E and F)** Gene Set Enrichment Analysis of unstimulated *MED12* deficient CAR-T cells compared to control cells using the Hallmarks gene collection. Normalized Enrichment Scores (NES) and FDR q values are shown. A positive NES indicates the gene set was enriched in *MED12* deficient cells. **G)** Chromatin accessibility profiles of *BATF3, IFNG, TNF, CCL3*, and *TOX* gene loci in control and *MED12* deficient CD19-28ζ CAR-T cells 15 days after T cell activation. **A, B, D, E, F)** Pooled data from *n* = 3 donors **C, G)** Representative donor of *n* = 3 donors.

Consistent with *MED12* deficient cells manifesting enhanced cytokine secretion and metabolic fitness, gene set enrichment analysis of differentially expressed genes revealed enrichment of metabolic and cytokine-related gene sets (**Fig. 4E-F**). Increased chromatin accessibility was most prevalent in introns and intergenic regions (**Fig. S7C**) in proximity to genes that regulate T cell memory and effector function, including *TBX21, TOX*, and *BATF3*^42–44^, suggesting that these regions function as enhancers (**Fig. 4C, 4G, Fig. S7D**). Together, the results demonstrate that *MED12* deficient T cells manifest widespread changes in the transcriptome, proteome and functional state associated with enhanced chromatin accessibility, most notably at key genes regulating effector T cell differentiation.

### Loss of MED12 increases core Mediator chromatin occupancy at transcriptionally active enhancers

The Mediator complex lacks a DNA binding domain, but interacts with chromatin through protein-protein interactions with DNA-bound transcription factors and RNA polymerase II (RNAPII)^41^. To identify the genomic locations of chromatin-Mediator interactions, we performed ChIP-seq using antibodies against MED12 and MED1 to profile chromatin binding of the kinase module and core Mediator, respectively, in the presence or absence of MED12. Comparison of MED1- and MED12 bound genomic regions in control CD19-28ζ CAR-T cells showed MED1/MED12 co-localization in 86.1% of sites, while MED1 was found exclusively at 13.1% of sites and MED12 was found exclusively at only 0.7% of sites (**Fig. 5A**). These results demonstrate that, in human T cells, the CDK module rarely contacts chromatin in the absence of core Mediator and is present at the majority of sites occupied by the core Mediator complex, leading to the prediction that *MED12* deletion could have widespread effects on the function of the core Mediator.

**Figure 5.**
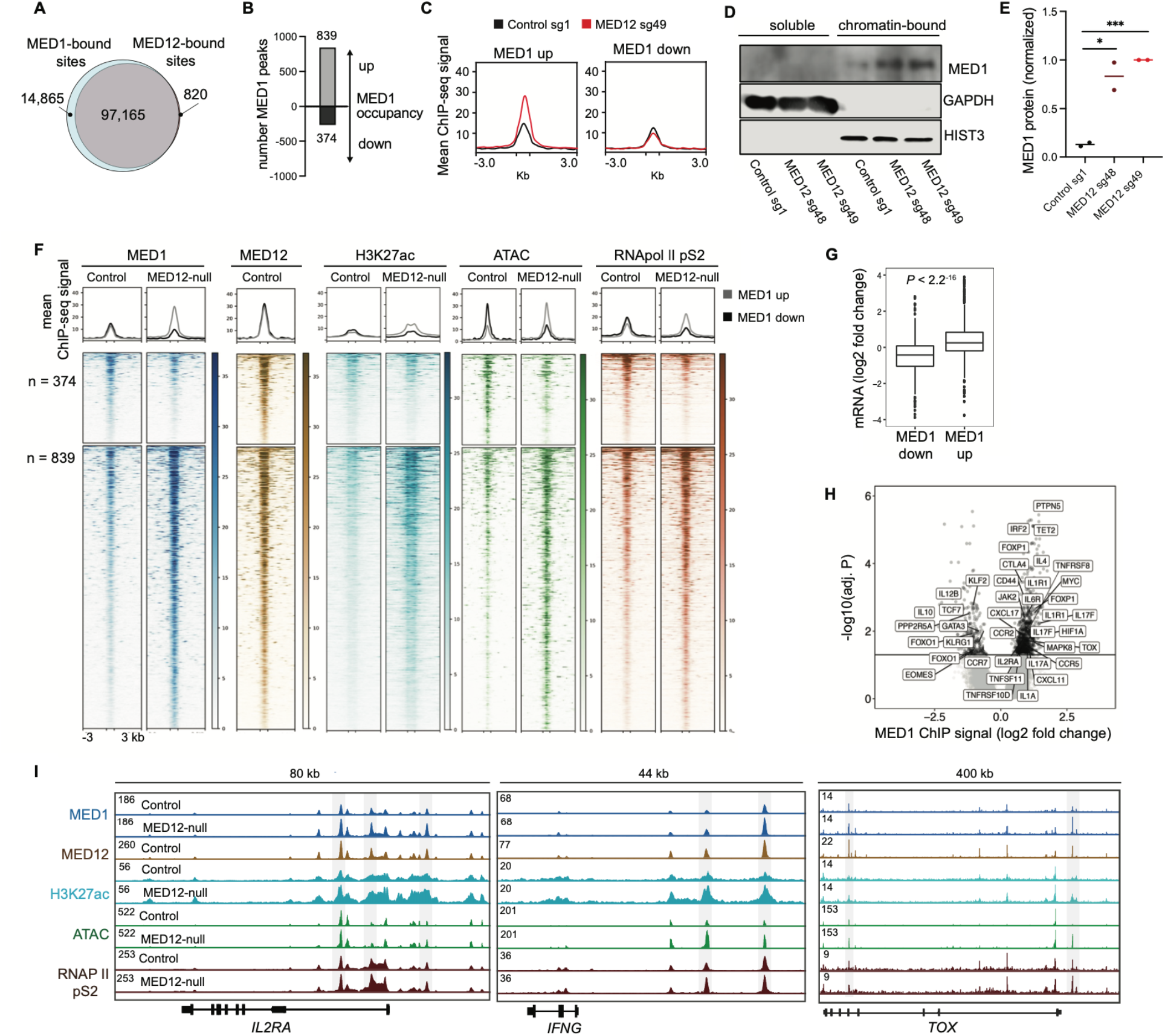
Loss of *MED12* increases MED1 chromatin occupancy at transcriptionally active enhancers. **A)** Venn diagram depicting number of sites bound by MED1 and/or *MED12* detected by ChIP-seq in CD19-28ζ control CAR-T cells. **B)** Number of genomic regions with significant change in MED1 occupancy detected by ChIP-seq between *MED12* deficient and control cells. Adjusted *P* < 0.05. **C)** Mean normalized ChIP-seq signal at regions with significant differences in MED1 occupancy. **D)** Western blot analysis of MED1 protein present in soluble and chromatin-bound cellular fractions from control and *MED12* deficient CD19-28ζ CAR-T cells 15 days after T cell activation. GAPDH and Histone 3 are used as markers for each cellular fraction. Representative blot from two independent experiments. **E)** Densitometric analysis of western blot shown in panel D. MED1 staining in the chromatin-bound fraction was normalized to HIST3 staining. *n* = 2 donors. Two-tailed unpaired Student’s *t*-test. \*\*\**P* < 0.001. **F)** Genomic regions with significantly different MED1 occupancy between Control and *MED12* deficient CD19-28ζ CAR-T cells overlaid with read coverage from *MED12*, MED1, RNAPII S2, and K3K27ac ChIP-seq and ATAC-seq. **G)** Box plot depicting fold change in mRNA expression between unstimulated *MED12* deficient and control CAR-T cells at genes proximal to genomic regions with differential MED1 occupancy. Kruskal Wallis test. **H)** Volcano plot depicting genes most proximal to genomic regions with differential MED1 occupancy between control and *MED12* deficient CD19-28ζ CAR-T cells (adjusted *P* < 0.05). **I)** ATAC-seq and ChIP-seq tracks at the *IFNG, IL2RA*, and *TOX* loci. **A, B, G, H)** Pooled data from *n* = 3 donors. **C, F, I)** One representative donor of *n* = 3 donors.

To assess the effect of MED12 deletion on core Mediator chromatin occupancy, we compared genomic regions bound by MED1 in control and MED12 deficient CAR-T cells. PCA showed global differences in MED1 occupancy between genotypes including 839 sites with increased MED1 occupancy and 374 sites with decreased MED1 occupancy (**Fig. S8A, Fig. 5B**), demonstrating a general pattern of increased MED1 binding in the absence of MED12. *MED12* deficient cells showed approximately a two-fold increase in MED1 ChIP-seq signal intensity at sites with significantly increased MED1 binding compared to control cells (**Fig. 5C**), a result that was confirmed by immunoblotting, since MED1 was more abundant in the chromatin-bound fraction in MED12 deficient cells compared with control cells (**Fig. 5D-E**). Furthermore, MED1 protein was more abundant in total cell lysates in *MED12* deficient CAR-T cells, while *MED1* transcript was not differentially expressed, indicating this effect was post-transcriptionally regulated (**Fig. S8B-D**).

To determine if sites with increased MED1 chromatin occupancy were transcriptionally active, we performed ChIP-seq with antibodies targeting H3K27ac and RNAPII pS2 (the elongating form of RNAPII). Comparison of peaks called at least once in any sample showed that 74.5% of sites displayed co-localization of MED1, MED12, H3K27ac, and RNAPII. Sites with increased MED1 binding also had increased H3K27ac, RNAPII occupancy, and chromatin accessibility (**Fig. 5F**). Furthermore, genes located in close proximity to sites with increased MED1 occupancy in *MED12* deficient cells correspond to significantly increased transcript and protein levels (**Fig. 5G**), including *IFNG, IL17F, IL2RA*, and *TOX* (**Fig. 5H-I**). Together, the data demonstrate widespread, but selective, enhancement of MED1 occupancy and transcriptional activation in *MED12* deficient CAR T cells, resulting in widespread transcriptional enhancement of genes regulating effector T cell differentiation, thereby leading to enhanced functional potency.

### Loss of MED12 increases STAT5 activity in CD19-28ζ CAR-T cells

To define the most differentially regulated transcriptional programs in *MED12* deficient cells, we used HOMER motif enrichment analysis to identify differentially accessible transcription factor binding motifs in *MED12* deficient vs. control cells. The top enriched motifs were STAT5 and STAT1, transcription factors that drive cytokine mediated gene expression **(Fig. 6A-B**). AP-1 family motifs including JunB, BATF, and FOS were also significantly enriched, as well as motifs from Interferon Response Family (IRF) members including IRF8 and IRF4. Motif enrichment analysis on sites with increased MED1 occupancy in MED12 deficient cells also showed enrichment of STAT and AP-1 motifs (**Fig. 6C**), confirming that core Mediator is recruited to these motifs.

**Figure 6.**
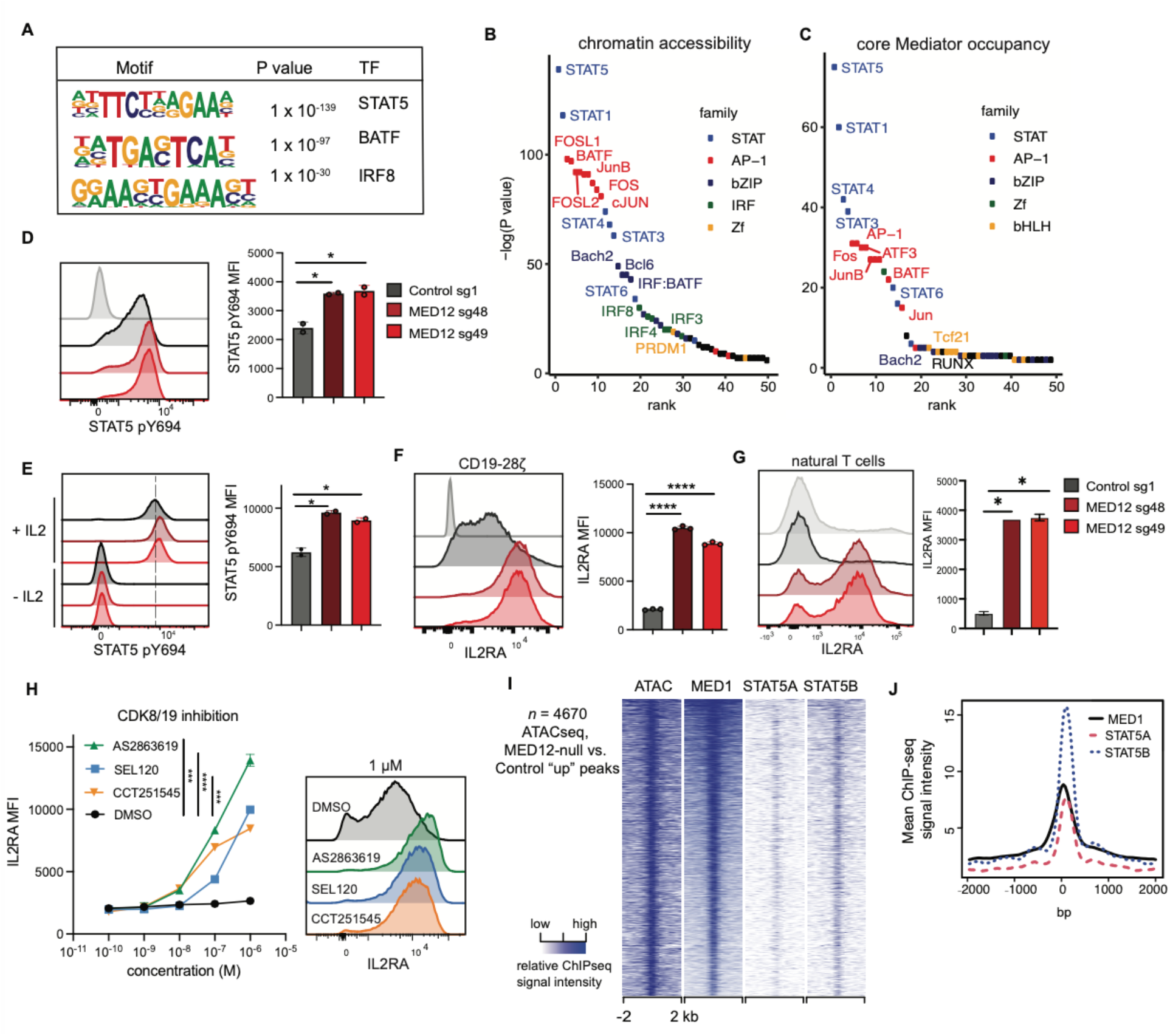
Loss of *MED12* increases STAT5 activity in CD19-28ζ CAR-T cells. **A and B)** Transcription factor binding motif enrichment at sites with increased chromatin accessibility by ATAC-seq in *MED12* deficient CD19-28ζ CAR-T cells 15 days post T cell activation. *n* = 4670 sites with Log2FC > 1 and adj. *P* < 0.05. **C)** Transcription factor binding motif enrichment at sites with increased MED1 occupancy by ChIP-seq in *MED12* deficient CD19-28ζ CAR-T cells. *n* = 839 sites with adjusted *P* < 0.05. **D)** Flow cytometry analysis of STAT5 phosphorylation in control and *MED12* deficient CD19-28ζ CAR-T cells 15 days post T cell activation. Cells were cultured *in vitro* with continuous IL-2. The cells shown in light grey were rested without IL-2 for 24 hours prior to staining. **E)** Flow cytometry analysis of STAT5 phosphorylation in control and *MED12* deficient CD19-28ζ 15 days post T cell activation. On day 14, CAR-T cells were rested for 24 hours without IL-2 and restimulated for 15 minutes with IL-2 prior to staining. **F and G)** Flow cytometry analysis of IL2RA expression in control and *MED12* deficient F) CD19-28ζ CAR-T cells or G) non-transduced T cells 15 days post activation. Control staining with isotype antibody is shown in light grey. **H)** Flow cytometry analysis of IL2RA expression in non-transduced T cells cultured in dual inhibitors of CDK8 and CDK19 for 15 days. Inhibitors were supplemented to the culture medium every 48 hours. **I)** Heatmaps of genomic loci with significantly increased chromatin accessibility by ATAC-seq in *MED12* deficient CD19-28ζ CAR-T cells. Genomic loci are overlaid with ATAC-seq and MED1 ChIP-seq signal from unstimulated CD19-28ζ CAR-T cells 15 days post T cell activation. STAT5A and STAT5B ChIP-seq data was obtained from human CD4^+^ T cells stimulated with IL-2 (GEO accessions GSM671400, GSM671402). **J)** Mean ChIP-seq signal intensities corresponding to panel G. **A-C)** Pooled data from *n* = 3 donors. Homer motif enrichment was performed with a set of all A and B) ATAC-seq or C) MED1 ChIP-seq peaks detected in CAR-T cells as the background. **D-H)** Data are mean ± s.d. from *n* = 2-3 wells. Representative results from two independent experiments. Two-tailed unpaired Student’s *t*-test. \**P* < 0.05, \*\**P* < 0.01, \*\*\**P* < 0.001, \*\*\*\**P* < 0.0001 **I and J)** One representative donor of *n* = 3 donors.

To assess the functional significance of these findings, we compared STAT5 phosphorylation in *MED12* deficient vs. control cells during *in vitro* culture with IL-2 and observed that *MED12* deficient cells manifest significantly higher levels of phosphorylated STAT5 (**Fig. 6D**). Importantly, *MED12* deficient cells rested for 24 hours without IL-2 did not demonstrate phosphorylated STAT5, confirming that STAT5 activation was dependent on IL-2, whereas a brief restimulation of rested cells with IL-2 demonstrated *MED12* deficient CAR-T cells reached higher maximum levels of phosphorylated STAT5 **(Fig. 6E**). *MED12* deficient CAR-T cells also manifested increased expression of the high-affinity IL2 receptor, IL2RA, a known downstream target of STAT5 mediated transcription^45^ (**Fig. 6F**). Non-transduced natural T cells also showed elevated IL2RA expression in the absence of *MED12*, indicating this effect was not dependent on CAR signaling (**Fig. 6G**). Furthermore, T cells cultured with CDK8/19 inhibitors also manifested elevated expression of IL2RA, demonstrating that inhibition of CDK8/19 kinase activity is sufficient to elicit increased sensitivity to IL-2 in human T cells (**Fig. 6H**).

To investigate interactions between Mediator and STAT5, we compared MED1 chromatin occupancy with previously published STAT5 ChIP-seq data from CD4^+^ T cells^46^ at sites that have increased chromatin accessibility in *MED12* deficient cells and found extensive co-localization of MED1 and STAT5 (**Fig. 6I-J**). Together, these results demonstrate augmented STAT5 activity in *MED12* deficient T cells, and are consistent with a model wherein *MED12* knock-out broadly reprograms T cell effector differentiation and function by modulating MED1 chromatin occupancy and thereby enhancing functionality of multiple transcription factors that control T cell fate during effector differentiation, including STAT5, as well as others such as AP-1, IRF, and other STATs (**Fig. 7**)^47–49^.

**Figure 7.**
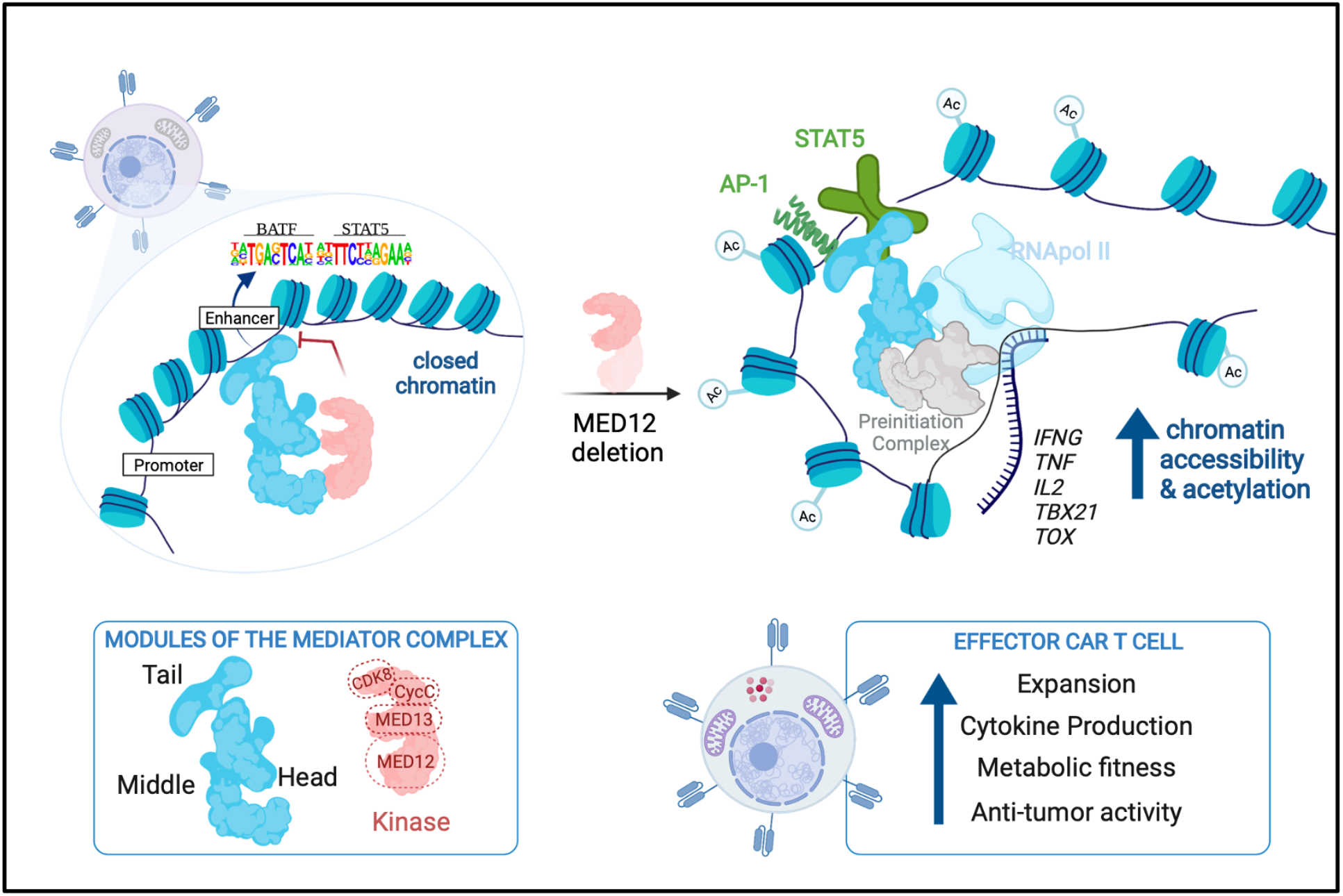
Proposed model of the mechanism by which the Mediator kinase module regulates T cell effector function. Disruption of the Mediator kinase module (pink) through loss of *MED12* leads to increased chromatin accessibility of STAT and AP-1 family transcription factor binding motifs, allowing core Mediator (blue) to associate with these regions. Chromatin accessibility and histone acetylation increases at these sites, thereby inducing transcription of T cell effector genes. Remodeling of the epigenetic state of CAR-T cells through loss of *MED12* promotes a sustained T cell effector phenotype characterized by increased expansion, metabolic activity, inflammatory cytokine production, and anti-tumor activity.

## Discussion

Immune checkpoint inhibitors and adoptive T cell therapies induce profound anti-tumor effects in some patients for whom all other therapies have failed, but most patients treated with cancer immunotherapies do not experience long-term benefit^1–5^. Enhancing the efficacy of cancer immunotherapies requires new approaches to augment the potency of tumor-specific T cell responses. In the context of adoptive cell therapy, significant efforts have focused on enhancing stem-like pools of memory T cells and thereby increasing the supply of effector cells^50–52^ and on preventing or reversing T cell exhaustion to enhance T cell function^13–15^. Here, we used systematic, unbiased genome-wide CRISPR-Cas9 screens to identify targets capable of enhancing CAR-T cell effector function. Our results converged on *MED12* and *CCNC*, core components of the Mediator kinase module, which have not previously been implicated in regulating T cell potency. Our screen also confirmed an expected requisite role for core Mediator in mature T cell function. *MED12* and *CCNC* deficient CAR-T cells demonstrate enhanced antigen-driven expansion, inflammatory cytokine secretion, cytolytic capacity, metabolic fitness and anti-tumor effects. The enhancements are observed in exhausted and non-exhausted T cells, in CAR-T cells targeting three distinct antigens and expressing 4-1BB or CD28 costimulatory domains, highlighting the generalizability of the findings.

Our data demonstrate that the CDK module and core Mediator are largely co-localized in wildtype CAR-T cells, consistent with the known role for the kinase module in modulating the interaction between core Mediator and RNAPII^53,54^, and leading to the hypothesis that loss of *MED12* or *CCNC* in T cells reduces steric hindrance between core Mediator and RNAPII, and thereby increases transcription and modulates T cell differentiation. Deletion of *MED12* shared some phenotypic similarities with deletion of *CCNC*, although more profound effects were observed following *MED12* deletion, consistent with the structural relationships that predict full ablation of the Mediator kinase module in the absence of *MED12*, and partial ablation upon *CCNC* deletion^29,53,55,56^. Our data demonstrate widespread increases in chromatin accessibility and enhancement of MED1 binding to chromatin in *MED12* deficient T cells. Further, we observed that *MED12* deletion induces genome-wide perturbations involving many transcription factors, thus explaining why the Mediator kinase module was the top hit in our CRISPR screens, rather than any single transcription factor.

While the effects of *MED12* deletion were widespread, they selectively enhanced expression of numerous transcription factors involved in T cell effector differentiation, including TOX, T-bet, and BATF3. Similarly, loss of *MED12* orchestrated coordinated changes in chromatin accessibility and increased MED1 occupancy at motifs for several transcription factor families involved in T cell differentiation including STAT5, STAT1, STAT4, STAT3 and Fos, cJUN, BATF, IRF8 and ATF3. The selectivity of the effects may be explained by the propensity for Mediator to augment transcription of genes regulated by super-enhancers, as described in setting of acute myelogenous leukemia, wherein inhibition of Mediator kinases selectively increased the activity of super-enhancer regulated transcription factors that program cell identity and cell fate^57^. In this regard, we interpret the effects of *MED12* and *CCNC* deletion to be highly context dependent, since we observed increased glycolytic rates in *MED12* and *CCNC* deficient effector T cells, consistent with enhanced effector activity, whereas previous studies in cancer have demonstrated diminished glycolytic activity in several cancers following small molecule mediated inhibition of CDK8^58^. Pharmacological inhibition of Mediator kinase activity phenocopied genetic ablation of *MED12* in regards to increased expansion and elevated expression of IL2RA, raising the possibility of synergistic antitumor effects in the context of immunotherapy since hyperactive CDK8/19 kinase activity activates oncogenes in some cancer types^59^.

Functional studies confirmed elevated STAT5 activity in *MED12* deficient T cells, manifested as increased sensitivity of T cells to IL-2. We also observed significant alterations in effector T cell differentiation. Current models hold that upon antigen encounter, naïve T cells differentiate into CCR7^−^IL7R^−^Tbet^+^CD62L^−^ short-lived effector cells (SLECs), with most effectors progressing toward a state of terminal differentiation associated with diminished activity, while a fraction express CCR7 and IL7R and differentiate into long-lived memory cells^36,38,60,61^. The functionally enhanced *MED12* deficient effector CAR-T cells observed here displayed a unique CCR7^−^IL7R^−^ Tbet^+^CD62L^+^ phenotype, with diminished expression of CD45RA, suggesting a block or delay in terminal effector differentiation^34^. While HA-28*ζ MED12* and *CCNC* deficient cells showed increases in LAG3, PD1 and TIM3 expression, which have been associated with T cell exhaustion, these markers are also associated with T cell activation. We did not observe upregulation of CD39 and observed augmentation of numerous functional attributes spanning improved proliferation, cytokine secretion, and ultimately, improved tumor control *in vivo*. Together these data are not consistent with exacerbation of T cell exhaustion in *MED12* deficient T cells.

In summary, this work identifies the Mediator CDK module as a critical negative regulator of T cell effector differentiation and function. Deletion of either *MED12* or *CCNC* induces broad based functional enhancements in effector T cells, including increased expansion, cytokine secretion, metabolic fitness and antitumor activity and phenotypic evidence of diminished terminal effector differentiation, all properties that would be predicted to enhance antitumor effects. Functional enhancements are also observed using small molecule mediated inhibition of CDK8/19. The work further therefore implicates, for the first time, interactions between the CDK module and core mediator as a major axis of regulation of T cell differentiation. Technologies to inactivate genes in the context of *ex vivo* cell manufacturing and even *in vivo* gene editing^17^, using a variety of approaches including CRISPR/Cas9, Zinc finger nucleases, TALENs, or base editing, are increasingly available for emerging applications in human medicine^62^, highlighting the potential for clinical translation of these findings.

## Acknowledgement

This work was supported by:

National Institutes of Health U54CA232568-01 (C.L.M.)

RM1-HG007735 (H.Y.C.)

R35-CA209919 (H.Y.C.)

Stand Up 2 Cancer–St. Baldrick’s–National Cancer Institute Pediatric Cancer Dream Team Translational Research Grant (SU2CAACR-DT1113, C.L.M.)

Parker Institute for Cancer Immunotherapy (A.T.S., H.Y.C., and C.L.M.)

Virginia and D.K. Ludwig Fund for Cancer Research (C.L.M.)

Sponsored research award from Lyell Immunopharma (C.L.M.)

Stand Up 2 Cancer is a program of the Entertainment Industry Foundation administered by the American Association for Cancer Research. C.L.M., H.Y.C., and A.T.S. are members of the Parker Institute for Cancer Immunotherapy, which supports the Stanford University Cancer Immunotherapy Program. H.Y.C. is an Investigator of the Howard Hughes Medical Institute.

E.W.W. is a member of the Parker Institute for Cancer Immunotherapy and was funded by a Parker Institute Bridge Scholar Award.

R.G.M. is the Taube Distinguished Scholar for Pediatric Immunotherapy at Stanford University School of Medicine.

J.A.B was supported by a Stanford Graduate Fellowship and the National Science Foundation Graduate Research Fellowship under Grant No. DGE-1656518.

K.A.F. was supported by the National Science Foundation Graduate Research Fellowship under Grant No. DGE-1656518.

## Author contributions

Conceptualization: CLM, ES, ATS, RGM, EWW, KAF,

Methodology: KAF, JAB, BD, KS, DK, PX, EWW, RGM, MM, ATS

Investigation: KAF, JAB, BD, KS, DK, PX, MM, VTD, ES, KAB, MGD

Visualization: KAF, JAB, VTD, KAB

Writing – original draft: KAF, JAB, ES, CLM

Writing – review & editing: HYC, ATS, EWW, BD, KS, JME

## Competing interests

K.A.F., E.S., and C.L.M. are coinventors on a patent for the use of T cells deficient in *MED12* or *CCNC* for therapeutic use. C.L.M. holds multiple patents in the arena of CAR T cell therapeutics.

C.L.M. is a cofounder and holds equity in Lyell Immunopharma and Syncopation Life Sciences, which are developing CAR-based therapies, and consults for Lyell, Syncopation, NeoImmune Tech, Apricity, Nektar, Immatics, Mammoth and Ensoma.

A.T.S. is a cofounder of Immunai and Cartography Bio.

H.Y.C. is an inventor on patents for the use of ATAC-seq, H.Y.C. is a co-founder of Accent Therapeutics, Boundless Bio, and an advisor for 10x Genomics, Arsenal Biosciences, Cartography Bio and Spring Discovery.

E.W.W. consults for and holds equity in Lyell Immunopharma and VISTAN Health.

E.S. consults for and holds equity in Lyell Immunopharma.

R.G.M. is a co-founder of and holds equity in Syncopation Life Sciences. RGM is a consultant for Lyell Immunopharma, Syncopation, Zai Lab, NKarta, and Aptorum Group.

J.A.B. is a consultant to Immunai.

## Data Availability

ATAC-seq, ChIP-seq, and RNA-seq data has been deposited into the Gene Expression Omnibus (GEO) repository under the accession number GSE174282. ATAC-seq and ChIP-seq data can be viewed through the UCSC Genome Browser at the following link: https://genome.ucsc.edu/s/kfreitas/med12%2Dcart%2Dv3

## Supplementary Materials

Materials and Methods

Fig. S1 to S8

Tables S1 to S5

