## Supplemental Materials for "Enhanced Effector Activity of Mediator Kinase Module Deficient CAR-T Cells"

**This PDF file includes:**

Materials and Methods
Table S4
Fig. S1 to S8

**Other Supplementary Materials for this manuscript include the following:**

**Table S1** CRISPR library sgRNA sequences and raw read counts
**Table S2** Results of MAGeCK analysis
**Table S3** Antibodies used in CyTOF panel
**Table S5** CAR sequences

### Materials and Methods

#### T cell isolation

Whole blood buffy coats were obtained from the Stanford Blood Center from healthy volunteers under 41 years. T cells were isolated using the RosetteSep Human T Cell Enrichment Cocktail (Stemcell Technologies). T cells were stored in CryoStor cell cryopreservation media CS10 (Sigma Aldrich*)* in liquid nitrogen.

#### CRISPR screen

*T Cell activation and culturing*
200 million T cells from each of two donors were thawed on day 0 and activated with CD3/CD28 Dynabeads (Invitrogen) at a ratio of three beads per T cell. Cells were cultured in AIM-V medium (Gibco) supplemented with 5% FBS, 10 mM HEPES, 1X penicillin-streptomycin-glutamine supplement (Gibco), and 10 ng/mL recombinant IL-2, (21.8 IU/mL) (Peprotech). Cells were maintained at a density between 0.5 and 2 million per mL in T175 flasks.

*Lentiviral transduction*
The complete Bassik Human CRISPR Knockout Library was obtained from Addgene and amplified with Endura ElectroCompetent Cells (Lucigen). LentiX cells (Takara) were plated on 150 mm plates coated with poly-D-lysine (Corning) and transfected with 18 µg REV, 18 µg GAG/POL, 7 µg VSVg, 15 µg library vector, 3.38 mL Opti-MEM (Gibco) and 135 µL Lipofectamine 2000 (Invitrogen) per plate. Media was changed 24 hours after transfection and supernatant was harvested 48 hours after transfection. Lentiviral supernatant was concentrated with Lenti-X Concentrator (Takara) and added to the T cell culture medium 2 days post activation. On day 3, cells were assessed for mCherry expression by flow cytometry to confirm the percentage of transduced cells was between 8 and 12%.

*CAS9 electroporation*
On day 3, 100 µl reactions were assembled with 10 million T cells, 30 µg Alt-R S.p. Cas9 Nuclease V3 (IDT), 90 µl P3 buffer (Lonza), and 7 µL Duplex Buffer (IDT). Cell were pulsed with protocol EO115 using the P3 Primary Cell 4D-Nucleofector Kit and *4D Nucleofector* System (Lonza). Cells were recovered immediately with warm media for 6 hours prior to transduction with CAR. The electroporation protocol was repeated on day 5.

*Retroviral Transduction*
Retroviral supernatant was produced as previously described[13]. Briefly, 293GP cells were plated on poly-D-lysine (Corning) coated plates and transfected with RD114 envelope and HA-28ζ CAR encoding plasmids using Lipofectamine 2000 (Invitrogen). Media was changed 24 hours after transfection and supernatant was harvested 48 and 72 hours after transfection. On days 3 and 4, tissue culture plates were coated with RetroNectin (Takara), blocked with 2% BSA for 5 minutes, and incubated with retroviral supernatant for 2 hours at 32˚C, 3200 rpm. T cells were added to virus coated plates at a density of $1 \times 10^6$ /mL. On day 5, CD28/CD3 Dynabeads (Invitrogen) were removed using magnetic separation. Cells were cultured with puromycin at 2.5 µg/mL from days 7 to 10 to eliminate cells which did not express a guide.

*Expansion Screen*
The CAR-T cells transduced with the sgRNA library was cultured in T175 flasks and were passaged every other day. On day 15, 100 million NALM6-GD2 cells were added to 100 million T cells and co-cultured to day 23. 50% of the culture volume was discarded at each passage.

On day 23, duplicate samples of 30 million cells were collected for genomic DNA extraction. The plasmid DNA encoding the lentiviral sgRNA library was used to approximate the relative abundance of sgRNAs at the start of the experiment.

*Cytokine Production Screen*
On day 15, duplicate samples of 30 million cells were harvested from the total population for genomic DNA extraction. 100 million CAR-T cells were co-cultured for 6 hours with 100 million NALM6-GD2-GFP tumor cells with eBioscience Monensin Solution (Invitrogen) in 200 mL medium without IL-2. Intracellular cytokine staining was performed using the Cytofix/Cytoperm Kit (BD). Cells were stained with antibodies purchased from Biolegend specific for CD4 (clone SK3), CD8 (clone SK1), TNF-α (clone MAb11), and IL-2 (clone MQ1-17H12), and fixable viability dye eFluor506 (eBioscience). Cell sorting was performed at the Stanford Shared FACS Facility on a FACSAria II equipped with a 70μm nozzle. The top 10% of $TNF\alpha^+$ and $IL\text{-}2^+$ were sorted using individual gates for $CD4^+$ and $CD8^+$ cells. $CD4^+$ and $CD8^+$ cytokine high cells were pooled. A single sorted sample of approximately 5 million $TNF\alpha^+$ $IL\text{-}2^+$ cells were collected from each donor for genomic DNA extraction.

*Genomic DNA extraction and sequencing library preparation*
Technical duplicates were performed for genomic DNA extraction, library preparation, and sequencing. Genomic DNA was extracted from cell pellets using overnight lysis in SDS with proteinase K at 37˚C as previously described [63]. Briefly, protein was precipitated with ammonium acetate and genomic DNA was precipitated with isopropanol. All of the recovered genomic DNA was used as template for PCR to generate the sequencing libraries. The libraries were prepared as previously described [23]. Illumina sequencing adapters were added using custom primers and sequencing was performed on the Ilumina NovaSeq 6000 PE150 platform at a depth of $5 \times 10^7$ reads per sample. Sequencing was performed by Novogene (Sacramento, CA).

*CRISPR Screen Data analysis*
Guide sequences were extracted from FASTQ files and matched to the Bassik library index using a custom R script. Raw counts for each guide were provided as input to the MAGECK algorithm [24]. For the expansion screen, two replicates from plasmid DNA library were compared to 4 samples collected on day 23 (two from each donor). For the cytokine production screen, 4 samples collected on day 15 (two from each donor) were compared to 2 samples (1 from each donor) that were sorted for high cytokine expression. The MAGeCK algorithm was used to perform normalization, calculate log fold changes for guides and genes, and calculate adjusted *P* values.

**Targeted CRISPR gene editing**
Ribonucleoprotein (RNP) was preparing using synthetic sgRNA with 2'-O-methyl phosphorothioate modification (Synthego) diluted in TE buffer at 100 μM. 5 μl sgRNA was incubated with 2.5 μl Duplex Buffer (IDT) and 2.5 μg Alt-R S.p. Cas9 Nuclease V3 (IDT) for 30 minutes at room temperature. 100 μl reactions were assembled with 10 million T cells, 90 μl P3 buffer (Lonza), and 10 μl RNP. Cell were pulsed with protocol EO115 using the P3 Primary Cell 4D-Nucleofector Kit and *4D Nucleofector* System (Lonza). Cells were recovered immediately with warm media for 6 hours prior to transduction with CAR. Guides sequences: AAVS1-sg1 5' GGGGCCACUAGGGACAGGAU 3', CCNC-sg40 5' GAUGCCAAAAACACACAUGU 3', CCNC-sg46 5' GGAUUUAAAGUUUCUCUCAG 3', MED12-sg48 5' CCUGCCUCAGGAUGAACUGA 3', MED12-sg49 5' UAACCAGCCUGCUGUCUCUG 3'.

**Assessment of targeted CRISPR gene editing**

4-7 days after editing, genomic DNA was extracted with QuickExtract DNA Extraction Solution (Lucigen) and ~500 bp regions flanking the cut site were amplified with Phusion Hot Start Flex 2X Master Mix (New England Biolabs) according to manufacturer's instructions. Sanger sequencing traces were analyzed by Inference of CRISPR Editing (ICE) [30].

**Cell lines**

NALM-6 leukemia cells and 143B osteosarcoma cells were obtained from American Type Culture Collection. Cell lines were stably transduced with GFP and firefly luciferase. Nalm6-GD2 was engineered to stably express GD2 synthase and GD3 synthase to obtain surface expression of GD2 disialoganglioside. Single cell clones were selected for high expression of GFP, luciferase, and GD2. Cell lines were maintained in RPMI (Gibco) supplemented with 10 mM HEPES, 10% FBS, and 1X penicillin-streptomycin-glutamine supplement (Gibco).

**T cell expansion and viability assays**

T cells were activated for 4 days at a 1:3 ratio of T cells to anti-CD3/28 Dynabeads (Invitrogen). T cell expansion assays were performed with IL-2 in the culture medium at 10 ng/mL (21.8 IU/mL) unless indicated otherwise. Cell counts and viability measurements were obtained using the Cellaca Mx Automated Cell Counter (Nexcelom). Cells were stained with acridine orange and propidium iodide to assess viability. A portion of the culture volume was discarded at each passage and the fraction of cells maintained in culture was used to calculate total cell counts.

**CDK8 kinase inhibitor assays**

SEL120 (SEL120-34A) hydrochloride, AS2863619, and CCT251545 (Selleckchem) were reconstituted at 5 mM in DMSO and stored at -80˚C. Human primary T cells were plated in 96 well plates with 50,000 T cells per well. Inhibitors were added 24 hours after CD3/CD28 bead activation and were freshly supplemented every 48 hours. CD3/28 beads were removed on day 4 following activation. The reported IC50 for CDK8 is 4.4 nM, 0.6 nM, and 5 nM for SEL120, AS286319, and CCT251545 respectively. The IC50 for CDK19 is 10.4 nM, 4.3 nM, and 6.3 for SEL120, AS286319, and CCT251545 respectively.

**Serial stimulation assay**

Starting from 10 days post-activation, CAR-T cells were plated at a 1:1 ratio with $GFP^+$ tumor cells without IL-2. At 48 hours intervals, cell counts were recorded, and flow cytometry was performed to assess the ratio of T cells to tumor cells. Co-cultures were then collected and replated in fresh media and additional tumor cells were added to maintain a 1:1 effector to target ratio.

**Cytokine production assays**

$5 \times 10^4$ T cells and $5 \times 10^4$ tumor cells were co-cultured in 250 μl media without IL-2 in round bottom 96 well plates for 24 hours. Culture supernatants were collected and analyzed by ELISA. IL-2 and IFNγ were detected with the ELISA MAX kit (Biolegend) and TNF-alpha was detected with the Quantikine kit (R&D Systems). Bead-based multiplex cytokine detection assays were performed at the Human Immune Monitoring Center (Stanford University) using the Luminex platform.

**Incucyte cytotoxicity assay**

$5 \times 10^4$ tumor cells were co-cultured with variable numbers of CAR-T cells in 250 μl media without IL-2 in flat bottom 96 well plates for 80 hours. Time lapse microscopy images were obtained with the Incucyte Live Cell Analysis system (Essen Bioscience) at 10X magnification. Total green object integrated intensity (GCU x $\mu m^2$/image) was used to assess tumor killing. Effector to target cell ratios are indicated in the Figures or Figure legends.

**RT-qPCR**
RNA was extracted with RNeasy kit (Qiagen) and cDNA was synthesized with High-Capacity cDNA Reverse Transcription kit (ThermoFisher). RT-qPCR was performed with PowerUp SYBR Green Master Mix (ThermoFisher) using the Bio-Rad CFX thermocycler and CFX Manager software. Target gene expression was normalized to the 18S housekeeping gene using the 2ΔΔCt method. IFNG-F 5' TGACCAGAGCATCCAAAAGA 3', IFNG-R 5' CTCTTCGACCTCGAAACAGC 3', IL2-F 5' TGCATTGCACTAAGTCTTGCAC 3', IL2-R 5' AGTTCTGTGGCCTTCTTGGG 3', TNF-F 5' CACAGTGAAGTGCTGGCAAC 3', TNF-R 5' AGGAAGGCCTAAGGTCCACT 3', 18S-F 5' GCAGAATCCACGCCAGTACAAG 3', 18S-R 5' GCTTGTTGTCCAGACCATTGG 3'.

**Seahorse Assay**
Metabolic analysis was carried out using Seahorse Bioscience Analyzer XFe96. Briefly, $2 \times 10^6$ cells were resuspended in XF assay media supplemented with 25 mM glucose, 2mM glutamine and 1 mM sodium pyruvate and plated on a Cell-Tak (Corning)-coated microplate allowing the adhesion of CAR T cells. Mitochondrial stress and glycolytic parameters were measured *via* oxygen consumption rate (OCR) (pmol/min) and extracellular acidification rate (ECAR) (mpH/min), respectively, with use of real-time injections of oligomycin (1.5 μM), carbonyl cyanide-*4* (trifluoromethoxy) phenylhydrazone (FCCP; 1 μM) and rotenone and antimycin (both 1 μM). Respiratory parameters were calculated following manufacturer’s instructions (Seahorse Bioscience). All chemicals were purchased from Agilent unless stated otherwise.

**Flow Cytometry**
T cells were washed in FACS buffer (DPBS no calcium, no magnesium (Gibco) with 2% FBS). Cells were incubated on ice in FACS buffer with antibodies specific for cell surface markers for twenty minutes. STAT5 staining was performed with the Fix and Perm Cell Permeabilization kit (ThermoFisher) according to manufacturer's instructions. Antibodies used are in Table 1. Cells were stained with MitoTracker Green (Cell Signaling Technology) at 200 nM, 37˚C for 30 minutes. Cells were washed in FACS buffer and analyzed on a LSRFortessa (BD) with BD FACSDiva software.

**Western Blotting**
Total cell lysates were extracted in non-denaturing lysis buffer (150 mM NaCl, 50 mM Tris pH 8, 1% NP-40, 0.25% sodium deoxycholate with Halt Protease Inhibitor Cocktail (ThermoFisher Scientific). Chromatin-bound and soluble cellular fractions were prepared with cytoskeletal lysis buffer (10 mM PIPES-KOH (pH 6.8), 100 mM NaCl, 300 mm sucrose, 3 mM MgCl2, and Halt Protease Inhibitor Cocktail. Briefly, cells were washed in PBS, resuspended in lysis buffer, and incubated on ice for 20 minutes. Cells were centrifuged at 5,000 rpm to separate the soluble and chromatin bound fractions. The soluble fraction was cleared by centrifugation at 13,000 rpm. The chromatin-bound fraction was resuspended in Sample Reducing Buffer (Pierce) and incubated at 100˚C for 5 minutes. Protein concentration was assessed with the DC Protein Assay kit (Bio-Rad) and 20 μg total protein was loaded per sample. Equal volumes of soluble and chromatin fractions were loaded for each sample. SDS-PAGE electrophoresis was performed, and proteins were transferred to PVDF membranes for immunoblotting. Antibodies used are listed in Table 1. Immunofluorescence was detected with the Odyssey Imaging System (Licor), or chemiluminescence was detected with autoradiography film.

**Western Blot Quantification**

Images captured with autoradiography film were scanned at 600 dpi in 16-bit greyscale, and images captured with the Odyssey Imaging System were exported as JPEGs. Quantification was performed with ImageJ software. A region of interest (ROI) of equal size was used to measure the specific band and background signal in each lane. Pixel densities were subtracted from 255 to invert the image and the background values were subtracted from band values to adjust for background signal. For each sample, the background adjusted MED1 pixel density was divided by the same value from the HIST3 loading control to calculate a ratio of MED1 to HIST3. Ratios from donor 1 and donor 2 were normalized to the largest ratio collected in each independent experiment.

**Mice**

Immunocompromised NOD *scid* IL2Rgamma$^{null}$ (NSG) mice were purchased from JAX and bred in-house under sterile conditions. Mice were monitored daily by the Veterinary Service Center staff. Care and treatments were in compliance with Stanford University APLAC protocols. Leukemia cells and CAR-T cells were administered via intravenous injection. 143B osteosarcoma cells were administered by intramuscular injection. For some experiments, tumor burden was assessed prior to treatment and mice were randomized to groups to ensure equal tumor burden between treatment groups. Time of treatment and dosing is indicated in Figure legends. Researchers were blinded during administration of T cells. Leukemia progression was monitored using the Lago SII (Spectral Instruments Imaging). Quantification of bioluminescence was performed with Aura software (Spectral Instruments Imaging). Solid tumor progression was followed using caliper measurements of the injected leg area. Researchers were blinded to the treatment groups during solid tumor measurements. Mice were euthanized upon manifestation of paralysis, impaired mobility, poor body condition (score of BC2-), or when tumor diameter exceed 17 mm. Sample sizes of 5 mice per group were selected based on previous experience with these models. All experiments were repeated twice with different donors, and donors used for *in vivo* experiments were different from the screening experiments.

**Blood Analysis**

Blood was collected from the retro-orbital sinus into Microvette blood collection tubes with EDTA (Fisher Scientific). Whole blood was labeled with anti-CD45 (HI30, ThermoFisher) and red blood cells were lysed with FACS Lysing Solution (BD) according to manufacturer's instructions. Samples were mixed with CountBright Absolute Counting beads (ThermoFisher) prior to flow cytometry analysis.

**CyTOF sample preparation and data analysis**

$2 \times 10^6$ CAR-T cells were collected on day 15, washed in PBS, and resuspended in 250 nM cisplatin (Fluidigm) for 3 minutes. Cells were washed twice in cell staining medium (CSM, PBS with 0.05% BSA and 0.02% sodium azide) followed by fixing in 1.6% paraformaldehyde for 10 minutes at room temperature. Cells were washed twice in PBS, flash frozen on dry ice, and stored at -80˚C. Cells were thawed, washed in CSM, and barcoded with the Cell-ID 20-plex kit (Fluidigm) as previously described [64]. Samples were pooled and stained for cell surface markers for 30 minutes at room temperature. Intracellular staining was performed using Permeabilization buffer (eBioscience) for 45 minutes on ice followed by two CSM washes. Panel of antibodies can be found in Table 1. Cells were resuspended with Cell-ID Intercalator-ID (Fluidigm), washed twice in deionized water, resuspended in 1X EQ beads, and acquired on a Helios mass cytometer (Fluidigm). After acquisition, data was normalized using MATLAB-based software [65] and debarcoded using MATLAB-debarcoder. Fsc files were uploaded to the OMIQ platform for analysis (OMIQ.ai).

**Bulk RNA-seq**
CAR-T cells were collected on day 15 and processed without freezing. RNA was extracted using the RNeasy mini kit (Qiagen). RNA quality was assessed by BioAnalyzer (Agilent). Sequencing libraries were prepared by Novogene (Sacramento, CA) and 150 bp paired end sequencing at a depth of $3 \times 10^7$ reads per sample was obtained using the Illumina NovaSeq6000 platform. FASTQ files were generated by Novogene. Transcripts were quantified with Salmon and DESeq2 was used to identify differentially expressed genes. GSEA analysis was performed using GSEA software (Broad Institute).

**ChIP-seq**
ChIP-seq was performed as previously described with minor modifications [66]. CAR-T cells ($3 \times 10^6$) were double crosslinked by 50mM DSG (disuccinimidyl glutarate, #C1104 - ProteoChem) for 30 minutes followed by 10 minutes of 1% formaldehyde. Formaldehyde was quenched by the addition of glycine. Nuclei were isolated with ChIP lysis buffer (1% Triton x-100, 0.1% SDS, 150 mM NaCl, 1mM EDTA, and 20 mM Tris, pH 8.0). Nuclei were sheared with Covaris sonicator using the following setup: Fill level – 10, Duty Cycle – 5, PIP – 140, Cycles/Burst – 200, Time – 4 minutes). Sheared chromatin was immunoprecipitated overnight with the antibodies shown in Table 1. Antibody chromatin complexes were pulled down with Protein A magnetic beads and washed once in IP wash buffer I. (1% Triton, 0.1% SDS, 150 mM NaCl, 1 mM EDTA, 20 mM Tris, pH 8.0, and 0.1% NaDOC), twice in IP wash buffer II. (1% Triton, 0.1% SDS, 500 mM NaCl, 1 mM EDTA, 20 mM Tris, pH 8.0, and 0.1% NaDOC), once in IP wash buffer III. (0.25 M LiCl, 0.5% NP-40, 1mM EDTA, 20 mM Tris, pH 8.0, 0.5% NaDOC) and once in TE buffer (10 mM EDTA and 200 mM Tris, pH 8.0). DNA was eluted from the beads by vigorous shaking for 20 minutes in elution buffer (100mM $NaHCO_3$, 1% SDS). DNA was decrosslinked overnight at 65C and purified with MinElute PCR purification kit (Qiagen). DNA was quantified by Qubit and 10 ng DNA was used for sequencing library construction with the Ovation Ultralow Library System V2 (Tecan) using 12 PCR cycles. Libraries were sequenced on the Ilumina NovaSeq 6000 PE150 platform at a depth of $3 \times 10^7$ reads per sample.

**ATAC-seq**
Approximately 100k CAR-T cells were used from each sample. Nuclei were isolated with ATAC-LB (10mM Tris-HCl pH7.4, 10mM NaCl, 3mM MgCl2, 0.1% IGEPAL) and used for tagmentation using Nextera DNA Library Preparation Kit (Illumina) from three donors. After tagmentation DNA was purified with MinElute PCR Purification Kit (Qiagen). Tagmented DNA was then amplified with Phusion high-fidelity PCR master mix (NEB) using 14 PCR cycles. Amplified libraries were purified again with MinElute PCR Purification Kit. Fragment distribution of libraries was assessed with Agilent Bioanalyzer and libraries were sequenced on the Ilumina NovaSeq 6000 PE150 platform at a depth of $3 \times 10^7$ reads per sample. Sequencing was performed by Novogene (Sacramento, CA).

**ATAC-seq and ChIP-seq data processing**

*Quality control, aligning, and signal tracks*
Paired end fastq files were trimmed to remove adapters and low quality sequences using `fastp` and then were aligned to the hg38 reference genome using `hisat2` with the `--no-spliced-alignment` and `--very-sensitive` options. Duplicates were marked and removed with `picard MarkDuplicates.` Reads with high quality concordant alignments to human chromosomes chr1-chr22 and chrX (e.g., excluding chrY and chrM) were converted to BED files for downstream processing. BED files were normalized by using the number of fragments overlapping transcription start sites (TSS) genome wide, which controls for differences in sequencing depth and also library quality between samples. Normalized genome coverage tracks were exported

from R using `rtracklayer::export` and visualized using the Integrative Genomics Viewer (IGV). Plots were generated in R using ChIPpeakAnno (3.13)[67].

*Obtaining a union peak set and counts matrix*
Peaks were called for each replicate using MACS2. Reproducible peaks for each sample were determined as peaks present in at least 2 of 3 replicates and merged to create a disjoint peak set for each sample. Reproducible peaks for each sample were then merged into a disjoint union peak set encompassing all samples using our previously described iterative procedure which merges the peak sets by repeatedly removing less significant overlapping peaks until no overlaps remain. The peak by sample counts matrix was obtained by counting reads overlapping with each peak. The counts matrix was loaded into DESeq2 for differential analysis. Selected peak sets were also visualized across samples by using the bigwig track files described above with `plotHeatmap` from the `deepTools` analysis suite.

*Details of peak sets*
Differential "up" and "down" peak sets: obtained from DESeq2 pairwise comparisons using fdr >= 0.05 and sometimes an additional fold change threshold, as indicated. "Bound" peak sets: for each sample (where a sample is, for example "h3k27ac_MED12_unstim") bound peaks were determined based on thresholding the normalized counts matrix. The normalized counts matrix was obtained by multiplying each column (replicate) in the counts matrix by (1e6 / total reads in peaks in that replicate). The bound peaks for each sample were then determined as peaks which had at least 2 of 3 replicates with a normalized value greater than a threshold. A cutoff of 2 was selected empirically, which yielded between 60,000 and 120,000 peaks per sample. Finally, bound peak sets for each sample were merged into bound peak sets for each sample set (e.g., an overall h3k27ac CAR-T peak set) by taking the union.

*Motif analysis*
Motifs enriched in particular peak sets were analyzed using the HOMER `findMotifsGenome.pl` utility. Total bound peaks for each sample set were used as background, thus, the enrichments represent motif enrichments relative to general CAR-T specific peaks rather than enrichments relative to the whole genome.

Table S4: Antibodies used in flow cytometry, western blot, and ChIP-seq

| ANTIBODY | IDENTIFIER | SOURCE |
|---|---|---|
| FLOW CYTOMETRY | | |
| STAT5-pY694 | 47 | BD |
| CCR7 | 150503 | BD |
| PD-1 | J105 | eBioscience |
| LAG3 | 3DS22H | eBioscience |
| CD45RO | UCHL1 | eBioscience |
| CD45 | HI30 | eBioscience |
| CD39 | A1 | Biolegend |
| TIM3 | F38-2E2 | Biolegend |
| CD25 | 2A3 | BD |
| WESTERN BLOT | | |
| GAPDH | D4C6R | Cell Signaling |
| Histone-3 | 1B1B2 | Cell Signaling |

| | | |
|---|---|---|
| MED12 | 4529 | Cell Signaling |
| Cyclin C | Ab85927 | Abcam |
| MED1 | BLR037F | Bethyl |
| MED1 | sc5334 | Santa Cruz Biotechnology |
| Nucleolin | sc55486 | Santa Cruz Biotechnology |
| ChIP-seq | | |
| H3K27ac | Ab4729 | Abcam |
| MED1 | A300-793A | Bethyl |
| MED12 | A300-774A | Bethyl |
| RNAPIIpS2 | Ab5095 | Abcam |

### Fig. S1.

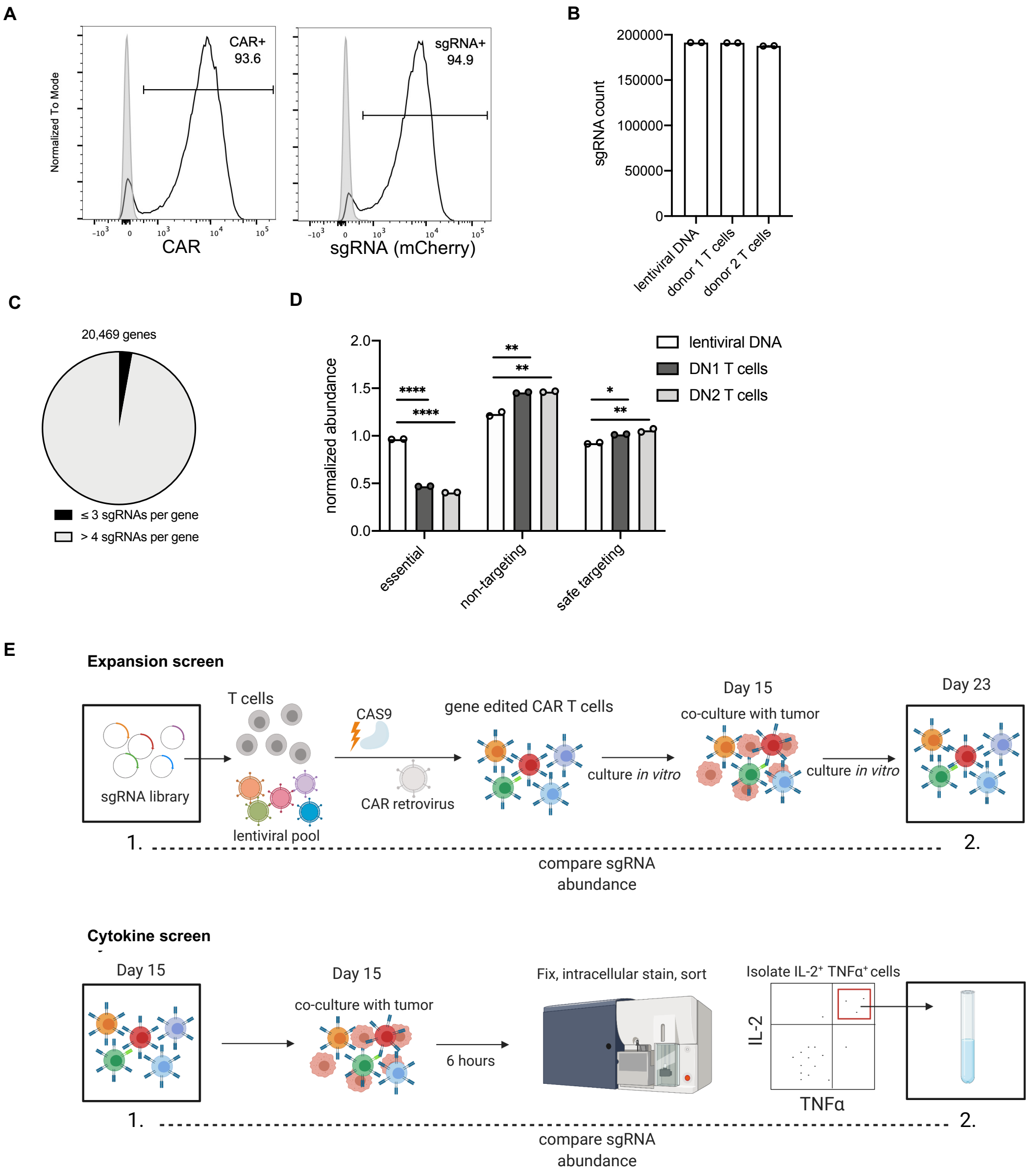


**Supplemental Figure 1. CRISPR screen library has adequate representation of guides and gene knockout is efficient.**
**A)** Flow cytometric histograms of the CAR-T cell library on day 15. (Left) HA-28ζ CAR expression detected by anti-idiotype antibodies. (Right) Integration of the sgRNA cassette detected by mCherry fluorescence.
**B)** Number of guides detected by deep sequencing of the lentiviral plasmid DNA library and CAR-T cell libraries on day 15.
**C)** Number of guides per gene detected in CAR-T cell libraries on day 15.
**D)** Abundance of guides targeting 214 essential genes, intergenic regions (safe targeting, $n$ = 6,750) and random sequence guides (non-targeting, $n$ = 5,642) in the lentiviral plasmid DNA library and day 15 CAR-T cell libraries. Two-tailed unpaired Student's $t$-test. *$P < 0.05$, **$P < 0.01$, ***$P < 0.001$, ****$P < 0.0001$
**B and D)** Data are mean ± s.d. of $n$ = 2 samples taken from the same pool.
**E**) Schematic depicting CRISPR screening for regulators of cytokine production (bottom) and T cell expansion (top). Box 1 and box 2 indicate which samples were compared to identify candidate genes.

### Fig. S2.

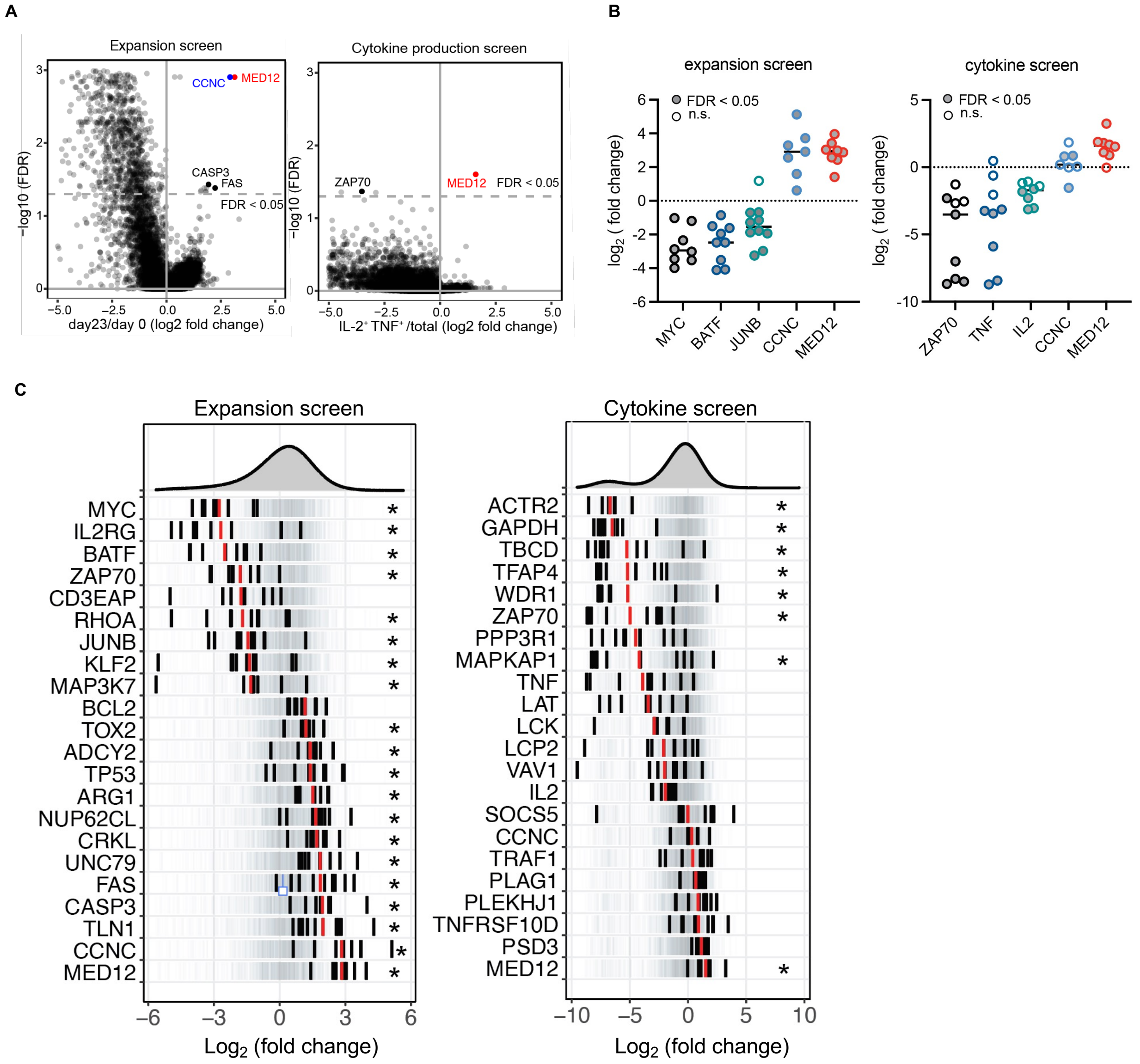


**Supplemental Figure 2. CRISPR screen identifies genes that are required for T cell survival and cytokine production.**

**A)** Volcano plot of gene knockout enrichment in expansion and cytokine production screens. (Left) Fold change in sgRNA abundance in the expansion screen. (Right) Fold change in sgRNA abundance in the cytokine production screen. The gene-level FDR was calculated using the MAGeCK algorithm.

**B)** Enrichment of individual sgRNAs. The sgRNA-level FDR was calculated using the MAGeCK algorithm. Significant sgRNAs are shaded in grey (FDR < 0.05).

**C)** Enrichment of sgRNA targeted selected genes. Individual sgRNAs are depicted as black lines and the mean as a red line. 1000 randomly selected guides are shown in light grey to indicate the library distribution. *Gene-level FDR < 0.05.

**A-C)** Data is pooled from *n* = 2 donors.

**Fig. S3.**

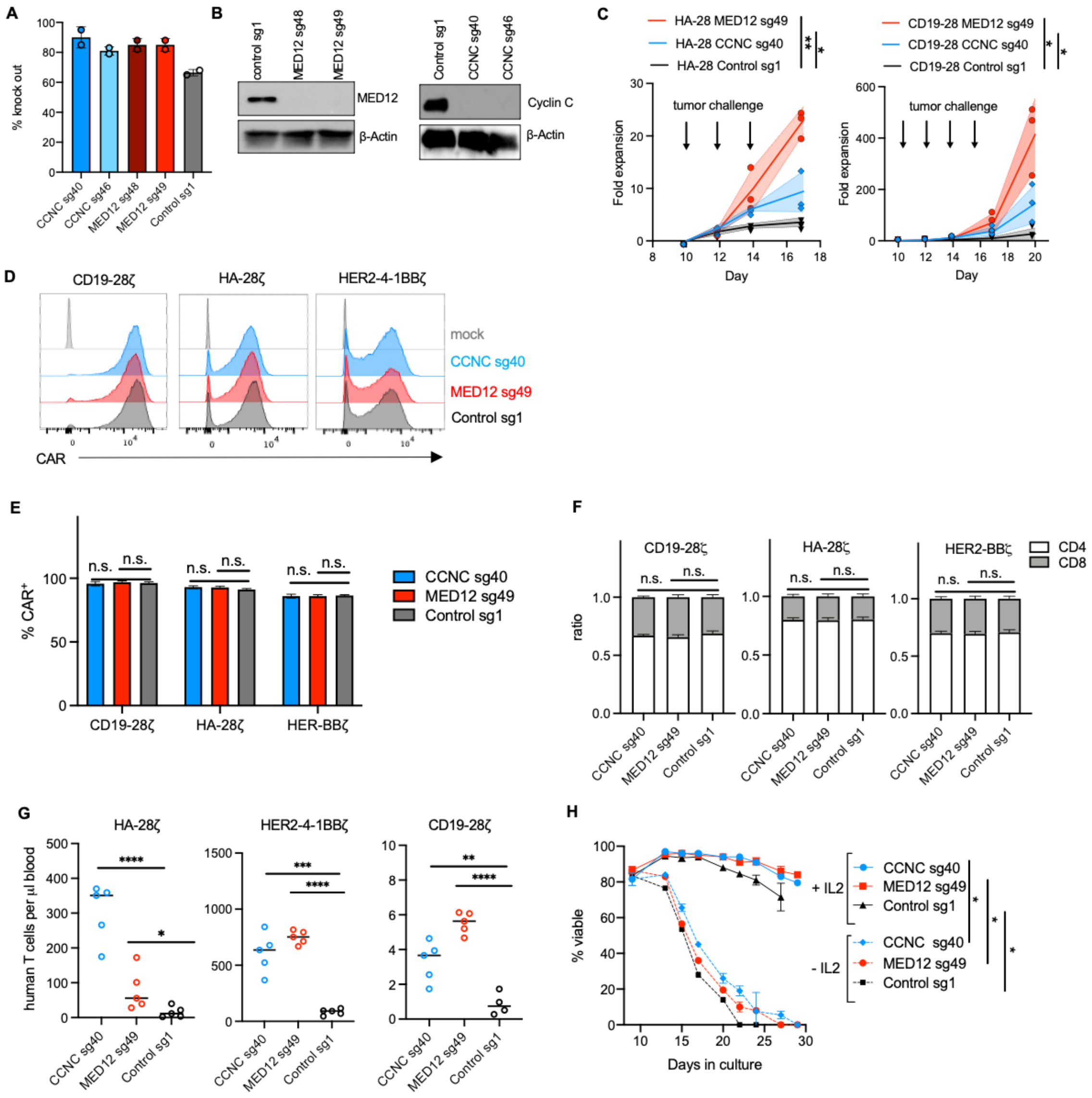

**Supplemental Figure 3. Target gene knockout and validation of expansion screen.**

**A)** Efficiency of target gene knockout inferred by Sanger sequencing and ICE analysis. Samples were collected 4 days after CRISPR editing. Mean ± s.d. (*n* = 2 donors).

**B)** Western blot confirming deletion of *CCNC* and *MED12*. Samples were collected 11 days after CRISPR editing. Representative blots of two independent experiments.

**C)** Antigen-driven *in vitro* expansion of *CCNC* and *MED12* deficient HA-28ζ (left) and CD19-28ζ (right) CAR-T cells. CAR-T cells were serially stimulated with $GD2^{+}$ or $CD19^{+}$ tumor cells in the absence of IL-2.

**D)** Flow cytometric histograms of surface CAR expression of control, *CCNC* or *MED12* deficient CD19-28ζ, HA-28ζ and HER2-4-1BBζ 10 days after T cell activation.

**E)** Percentage of $CAR^{+}$ T cells treated as in (C). Mean ± s.d. from *n* = 3 wells.

**F)** Ratio of $CD4^{+}$ to $CD8^{+}$ CAR-T cells treated as in (E). Mean ± s.d. from *n* = 3 wells.

**D - F)** Representative results of *n* = 4 donors (HA-28ζ, CD19-28ζ) or *n* = 2 donors (HER2-4-1BBζ).

**G)** Number of human T cells in peripheral blood from NSG mice bearing tumors shown in Fig. 2G-I 10 days post-infusion. Human T cells were detected by flow cytometric analysis of human CD45 expression. *n* = 5 mice. Results from three independent experiments are shown.

**H)** Cell viability of CD19-28ζ CAR-T cell with or without IL-2 supplemented to the culture medium. $n = 2$ replicate wells. Two-way ANOVA test with Dunnett's multiple comparison test. $^{*}P < 0.001$.
**E - G)** Two-tailed unpaired Student's $t$-test. $^{*}P < 0.05$, $^{**}P < 0.01$, $^{***}P < 0.001$, $^{****}P < 0.0001$

**Fig. S4.**

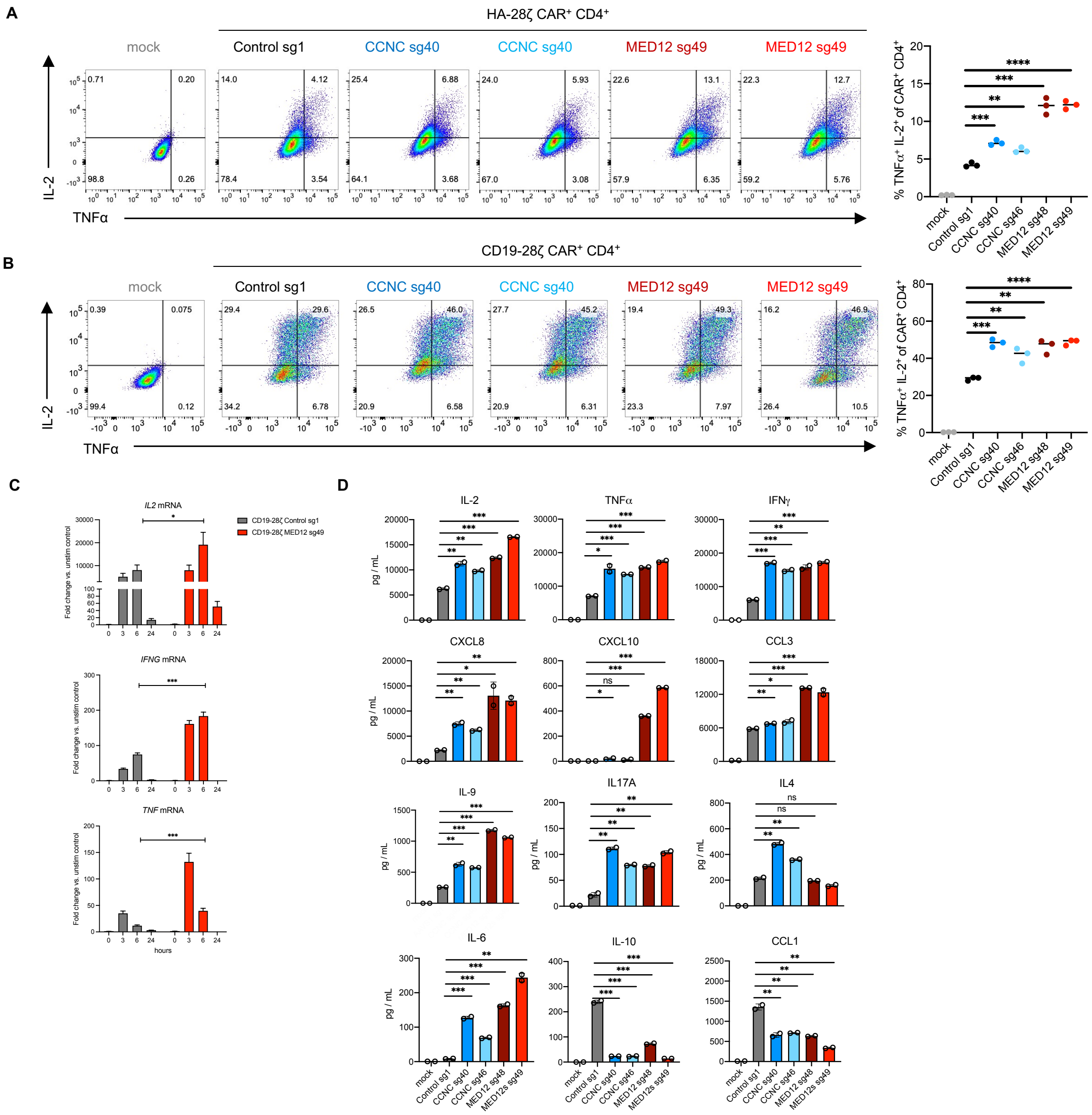


**Supplemental Figure 4. Validation of cytokine production screen.**

**A and B)** Flow cytometric dot-plots of intracellular IL-2 and TNFα on Control or *MED12* deficient CD19-28ζ CAR-T cells following 6-hour co-culture with tumor cells. CD19-28ζ and HA-28ζ were stimulated with NALM6 or NALM6-GD2 leukemia cells respectively. Bar graphs show percent of IL-2/TNF double positive CD4⁺ cells. Data are mean from triplicate wells. Representative result from *n* = 3 donors. Two unique sgRNAs were used to validate each candidate gene.

**C)** Relative mRNA expression of *IL2, IFNG* and *TNF* measured by RT-qPCR. Control or *MED12* deficient CD19-28ζ CAR-T cells were stimulated for 3, 6, or 24 hours with NALM6 leukemia cells. Mean ± s.d. (*n* = 3 wells).

**D)** Cytokines produced by Control or *MED12* deficient CD19-28ζ CAR-T cells following 24-hour co-culture with NALM6 leukemia cells measured by a multiplex bead-based assay. Data are mean ± s.d. from duplicate wells.

**A-D)** Two-tailed unpaired Student's *t*-test. All *MED12* deficient or *CCNC* deficient conditions were compared to the control. $^{*}P < 0.05$, $^{**}P < 0.01$, $^{***}P < 0.001$, $^{****}P < 0.0001$

**Fig. S5.**

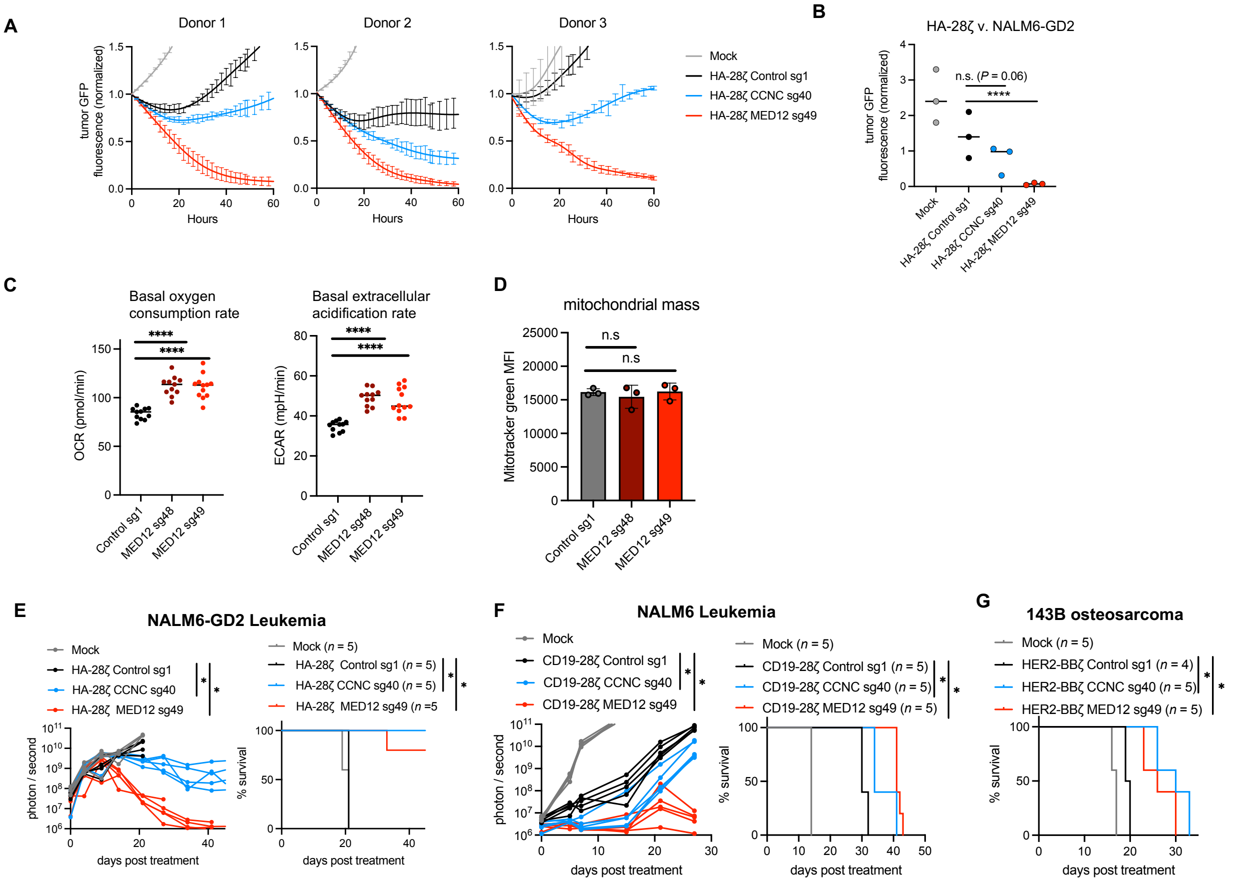


**Supplemental Figure 5. *MED12* deficient and *CCNC* deficient CAR-T cell have increased anti-tumor activity.**

**A)** Cytotoxicity of *CCNC* and *MED12* deficient HA-28ζ CAR-T cells against GFP$^+$ NALM6-GD2 leukemia following serial stimulation beginning 10 days after T cell activation. Cells were counted and replated at a 1:1 ratio of T cells to tumor cells at 48-hour intervals in media without IL-2. Data are mean ± s.d. from $n$ = 3 wells. Results from $n$ = 3 donors are shown.

**B)** Quantification of tumor fluorescence after 60-hours of co-culture shown in (A). $n$ = 3 donors**.** Ratio paired T test. ****$P$ < 0.0001.

**C)** Seahorse analysis of oxygen consumption rate (OCR) and extracellular acidification rate (ECAR) of control or *MED12* deficient CD19-28ζ CAR-T cells under resting conditions. Data are mean of $n$ = 12 replicate wells. Representative results from two independent experiments.

**D)** Flow cytometric analysis and quantification of mitochondrial mass in *CCNC* and *MED12* deficient CD19-28ζ CAR-T cells measured by Mitotracker Green 15 days after T cell activation ($n$ = 3 replicate wells). Representative result of two independent experiments.

**E and F)** Survival of CAR-treated mice shown in Fig. 2G, 2H. Survival curves were compared with the Log-rank Mantel-Cox test. *$P$ < 0.01. The number of mice per group is indicated in the Figure legend. Tumor growth was monitored by bioluminescent imaging. Two-way ANOVA test with Dunnett's multiple comparison test. *$P$ < 0.01

**G)** Survival of CAR-treated mice shown in Fig. 2I. Survival curves were compared with the Log-rank Mantel-Cox test. *$P$ < 0.01.

**E, F, and G)** Representative results of two independent experiments ($n$ = 2 donors).

**Fig. S6.**

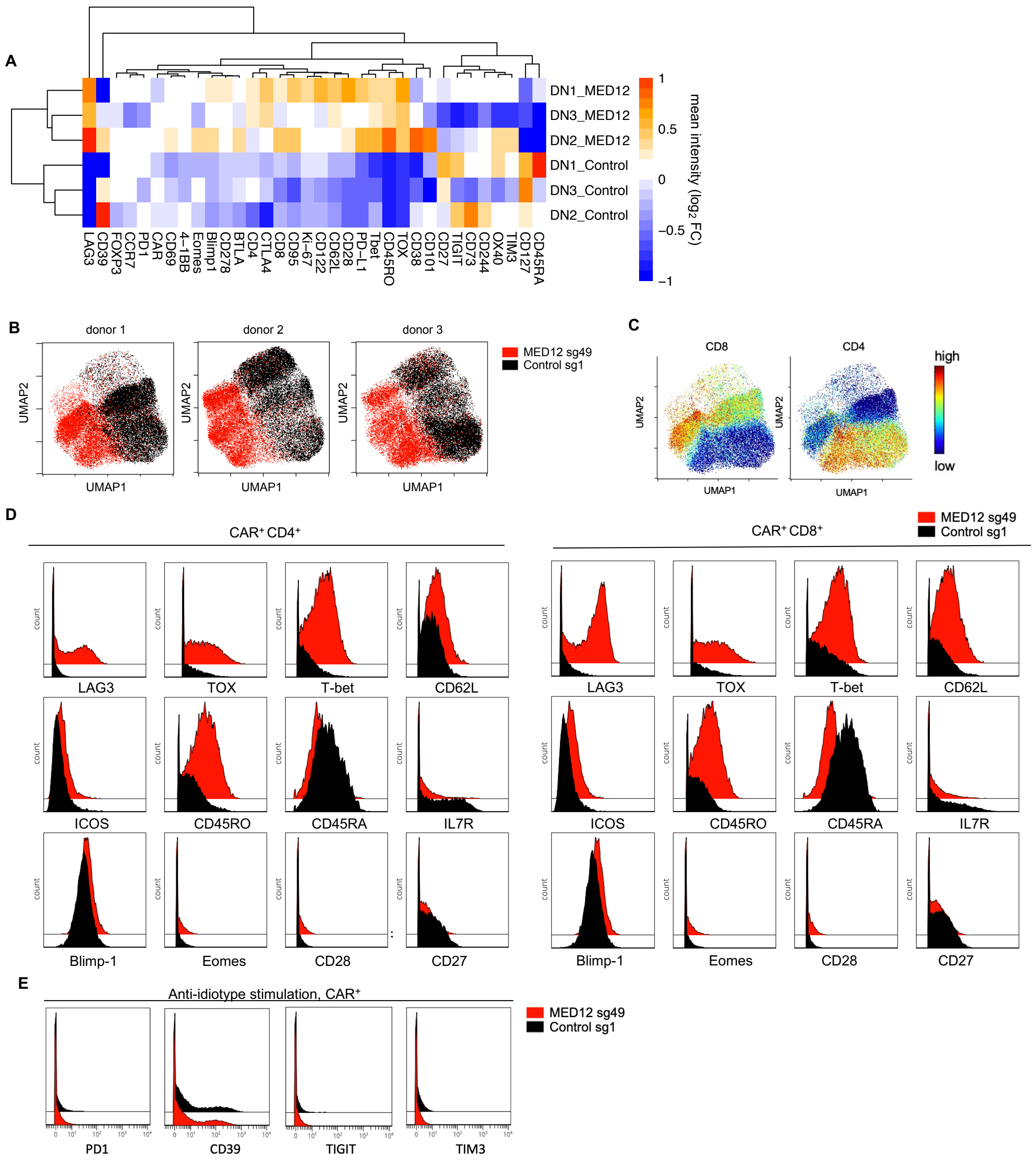


**Supplemental Figure 6. CyTOF analysis shows *MED12* deficient CAR-T cells have an effector-like phenotype.**

**A)** Heatmap depicting mean intensities of surface and intracellular markers analyzed by CyTOF in control or *MED12* deficient CAR$^+$ CD19-28ζ T cells 15 days after T cell activation. *n* = 3 donors.

**B)** Uniform Manifold Approximation and Projection (UMAP) analysis for each of the 3 donors. Control and *MED12* deficient files are overlaid and colored by genotype.

**C)** CyTOF analysis of CD4 and CD8 expression. Control and *MED12* deficient files are overlaid and colored by marker intensity (*n* = 3 donors).

**D and E)** Histograms displaying selected markers detected by CyTOF analysis. One representative donor is shown (*n* = 3 donors).

**A-D)** Cells were unstimulated.

**E)** Cells were stimulated through the CAR for 3 hours with anti-idiotype antibody.

### Fig. S7.

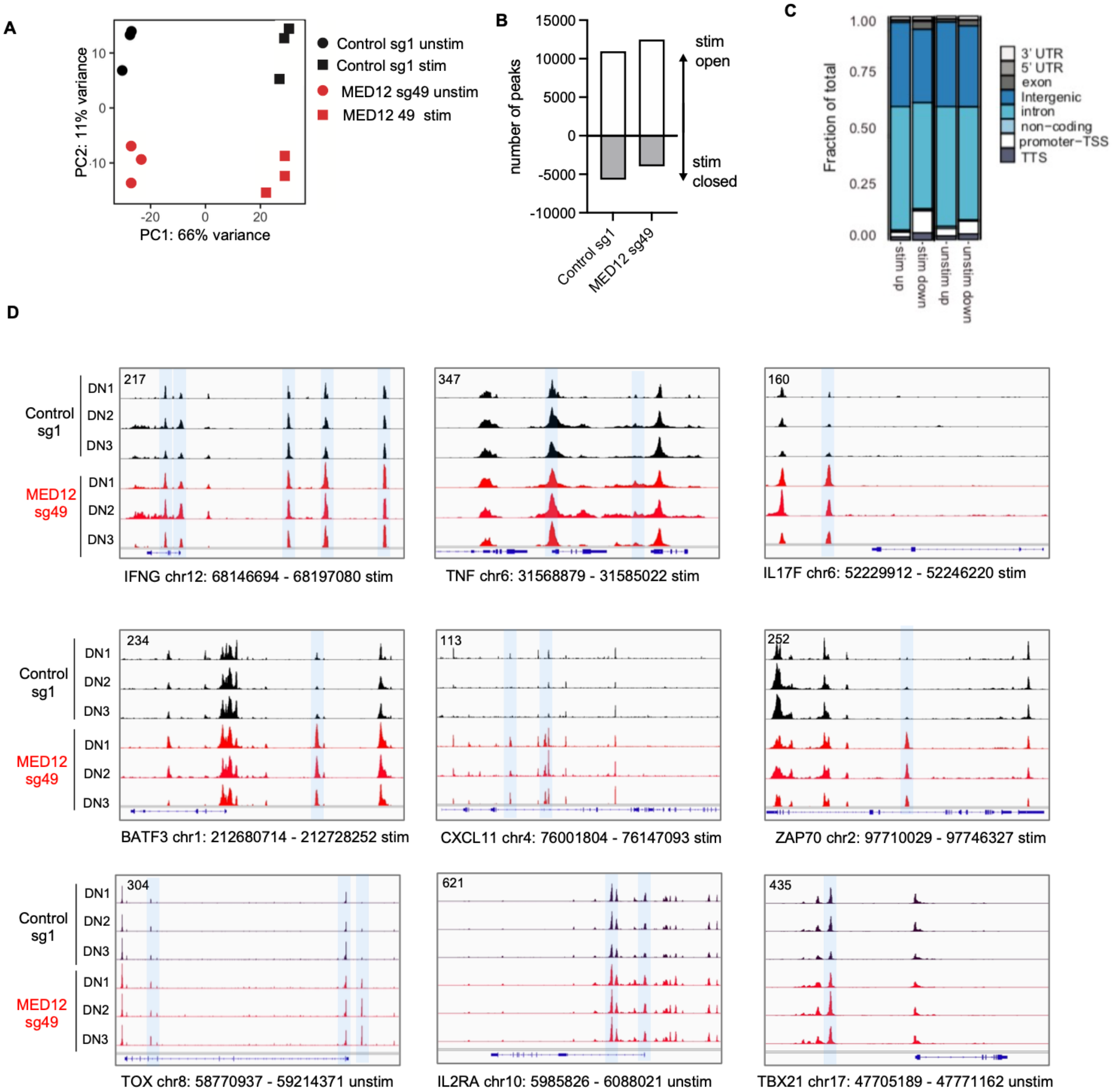


**Supplemental Figure 7. *MED12* deficient CD19-28ζ CAR-T cells have an effector-like transcriptomic profile and increased chromatin accessibility.**

**A)** PCA of bulk RNA-seq data from unstimulated (unstim) and stimulated (stim) control and *MED12* deficient CD19-28ζ CAR-T cells. Cells were cultured *in vitro* and stimulated with plate-bound anti-idiotype antibody for 3 hours prior to RNA extraction. *n* = 3 donors.

**B)** Number of peaks with significant change in accessibility detected by ATAC-seq between *MED12* deficient and control cells. Log2FC > 1 and adjusted $P < 0.05$.

**C)** Genomic annotations of ATAC-seq peaks that are differentially accessible between control and *MED12* deficient CD19-28ζ CAR-T cells.

**D)** ATAC-seq tracks of gene loci in stimulated (stim) and unstimulated (unstim) control or *MED12* deficient CD19-28ζ CAR-T cells on day 15. *n* = 3 donors.

### Fig. S8.

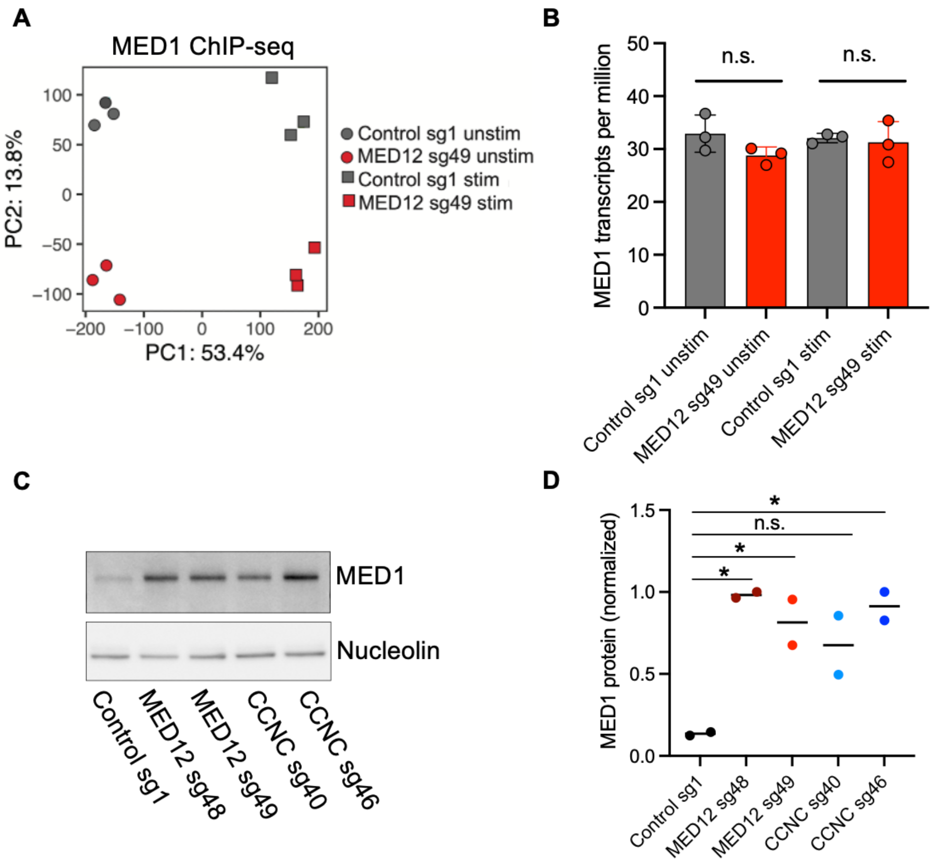


**Supplemental Figure 8. Increased levels of MED1 protein in MED12 deficient CD19-28ζ CAR-T is post-transcriptionally regulated.**

**A)** PCA of bulk MED1 ChIP-seq data from unstimulated (unstim) and stimulated (stim) control and *MED12* deficient CD19-28ζ CAR-T cells. Cells were cultured *in vitro* and stimulated with plate-bound anti-idiotype antibody for 3 hours prior to RNA extraction. *n* = 3 donors.

**B)** mRNA transcript levels from unstimulated (unstim) and stimulated (stim) control and *MED12* deficient CD19-28ζ CAR-T cells detected by RNA-seq. *n* = 3 donors. Statistical significance was determined by DESeq2.

**C)** Western blot analysis of MED1 protein present in total cell lysates from control and *MED12* deficient CD19-28ζ CAR-T cells 15 days after T cell activation. Representative blot from two independent experiments.

**D)** Densitometric analysis of western blot shown in panel C. *n* = 2 donors. Two-tailed paired ratio Student's *t*-test. **P* < 0.05.
